# Alcohol-specific transcriptional dynamics of memory reconsolidation

**DOI:** 10.1101/2022.06.09.495161

**Authors:** Koral Goltseker, Patricia Garay, Shigeki Iwase, Segev Barak

## Abstract

Relapse, a critical issue in alcohol addiction, can be attenuated by disruption of alcohol-associated memories. Memories are thought to temporarily destabilize upon retrieval during the reconsolidation process. Here, we characterized the alcohol-specific transcriptional dynamics that regulate these memories. Using a mouse place-conditioning procedure, we found that alcohol memory retrieval increased the expression of *Arc* and *Zif268* in the dorsal hippocampus (DH) and medial prefrontal cortex (mPFC). Alcohol seeking was abolished by post-retrieval non-specific inhibition of gene transcription in the DH, as well as by downregulating ARC expression in the DH using antisense-oligodeoxynucleotides. Since sucrose memory retrieval also increased *Arc* and *Zif268* expression, we performed an RNA-sequencing assay, and revealed alterations in the expression of *Adcy8, Neto1, Slc8a3* in the DH and *Fkbp5* in the mPFC, caused by the retrieval of alcohol but not sucrose memories. This offers a first insight into the unique transcriptional dynamics underpinning alcohol memory reconsolidation.

## Introduction

Alcohol use disorder (AUD) is a detrimental neuropsychiatric disorder with severe medical, social and economic burdens^1^, yet available pharmacotherapy is limited^2^. Nearly 70% of patients relapse within the first year of abstinence^3^, marking relapse as a major clinical challenge. Relapse is often triggered by craving for alcohol, evoked by environments and cues previously associated with alcohol^4^. Therefore, the disruption of memories that evoke alcohol-related behaviors is expected to reduce or even prevent cue-induced relapse^5, 6^.

It is increasingly accepted that well-consolidated memories can be reactivated upon retrieval. Retrieved memories undergo temporary destabilization and subsequent re-stabilization, a process termed reconsolidation^7-11^. Thus, memory reactivation initiates a 5-6 hour-long “reconsolidation window”, during which time a memory is labile for certain manipulations^7, 8, 11^. Indeed, interference with the reconsolidation of drug memories was shown to attenuate their subsequent expression and cue-induced relapse, thus providing a potential strategy for relapse prevention^12, 13^.

Although the exact mechanisms underpinning the processing of reactivated drug memories have yet to be characterized, reconsolidation of drug and alcohol memories were generally shown to be interrupted by the inhibition of NMDA^14-16^ or beta-adrenergic receptors^16, 17^; or by preventing protein synthesis^5, 10, 14^. According to recent fear and drug memory studies, memory reconsolidation requires not only protein synthesis but also gene transcription^18^. Moreover, the transcription of certain immediate early genes (IEGs), including *Arc*, encoding activity-regulated cytoskeleton-associated protein and the transcription factor-encoding *Zif268/Egr1*, was implicated in the reconsolidation of various types of memory^18-21^, implying that similar dynamics might control the reconsolidation of alcohol memories.

Our previous findings suggested that alcohol memory reconsolidation are processed via unique molecular mechanisms, not shared with non-alcohol memories. Specifically, we previously showed that inhibition of mechanistic target of rapamycin complex 1 (mTORC1), which controls the synthesis of a subset of dendritic proteins^22^, disrupted the reconsolidation of alcohol memories but not of memories associated with sucrose^5^. Therefore, it is possible that alcohol memory reconsolidation is characterized by a unique transcriptional profile. As such, we sought to determine the transcriptional dynamics that underlie alcohol memory reconsolidation within the dorsal hippocampus (DH) and medial prefrontal cortex (mPFC)^5, 23, 24^, brain regions implicated in alcohol use disorder^25, 26^ and in the formation, retention and expression of drug memories^5, 27, 28^.

## Results

### Alcohol memory reconsolidation depends on *de novo* gene transcription in the DH

While it has been established that the reconsolidation of alcohol memories requires *de novo* protein synthesis^5, 14^, it remains unclear whether it is also dependent on *de novo* gene transcription. Therefore, we assessed the role of gene transcription during alcohol memory reconsolidation within the DH, a brain region implicated in alcohol use disorder^25^ and involved in drug memory formation, retention, and expression^27, 28^, in addition to memory reconsolidation^29, 30^. To form alcohol-associated memories, we employed the alcohol-conditioned place preference (CPP) paradigm. This paradigm has been used to examine the reinforcing properties of alcohol, as well as to explore the processing and maintenance of memories that evoke relapse to alcohol-seeking in rodents^31, 32^, particularly in the DH^33^.

To assess the role of hippocampal gene transcription in alcohol memory reconsolidation, we formed alcohol-associated memories in the alcohol-CPP procedure, by conditioning one compartment of the CPP-box to alcohol (Figure 1A, experimental design). A day after confirming the strong preference for the alcohol-paired compartment in a CPP test, the mice were re-exposed to the alcohol-paired compartment for 3 min to retrieve alcohol-associated memories, as we previously demonstrated^34, 35^. Immediately after memory retrieval, actinomycin (4 µg/µl; 0.5 µl per side) or vehicle were infused into the DH^18^. In a retention test conducted 24 h later, we found that mice that received post-retrieval actinomycin D did not show alcohol-CPP, whereas the preference for the alcohol–associated compartment remained high in the vehicle-treated mice (Figure 1B; see Figure S1 for individual data). Thus, inhibition of gene transcription in the DH following memory retrieval led to a loss of alcohol-CPP, suggesting that the alcohol memory reconsolidation requires *de novo* gene transcription in the DH.

**Figure 1.**
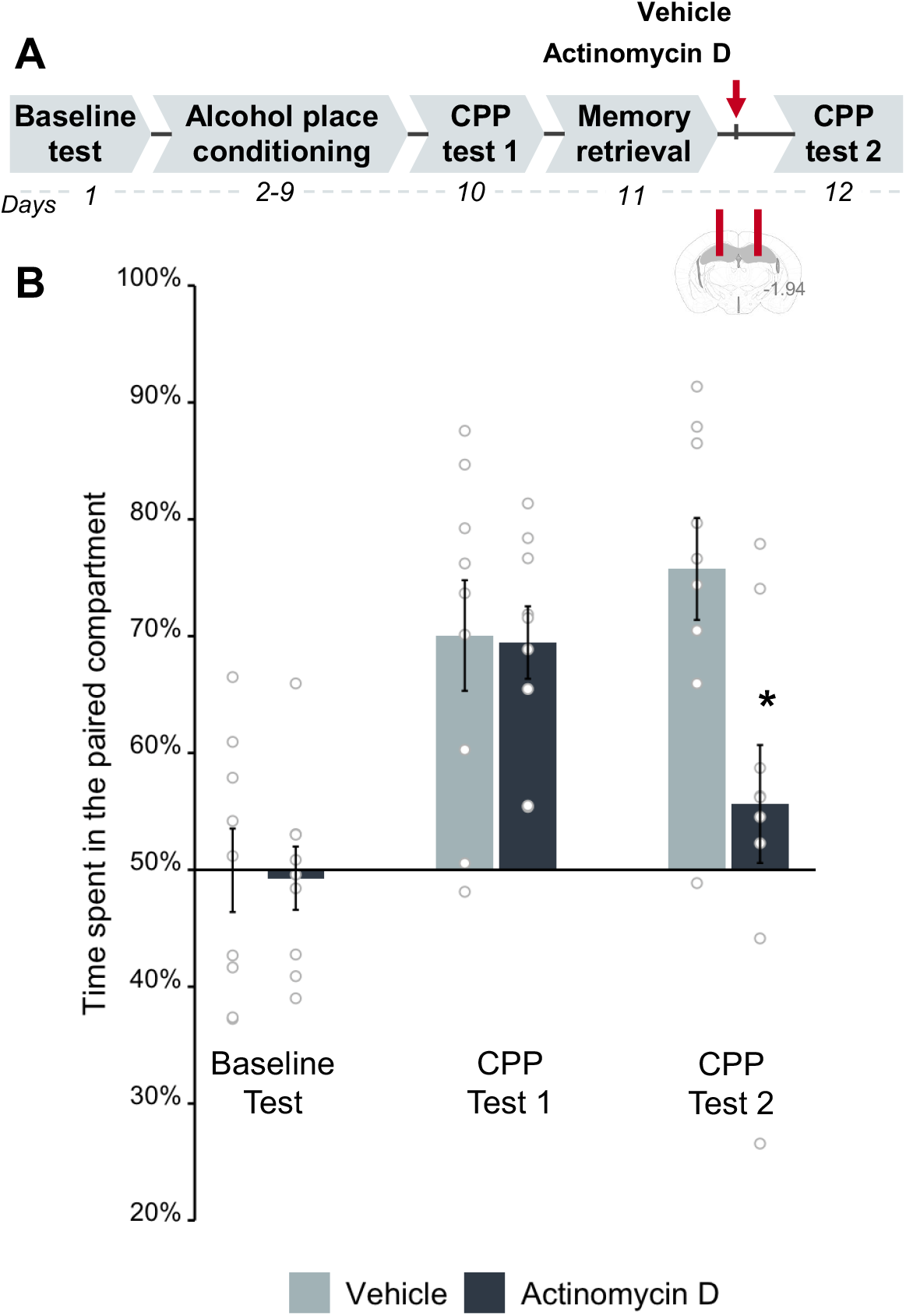
Inhibition of transcription in the dorsal hippocampus after alcohol memory retrieval disrupts the expression of alcohol-conditioned place preference (CPP). **A**. Schematic illustration of the experimental design and timeline. Actinomycin D (4 µg/µl) was bilaterally infused into the dorsal hippocampus of mice immediately following the retrieval of alcohol memories. **B**. Place preference scores, expressed as means ±S.E.M. of the percent of time spent in the alcohol-paired compartment. Mice that showed strong alcohol-CPP (t^(17)^=8.31, p<0.0001) lost alcohol-place preference when memory retrieval was followed by intra-DH infusion of actinomycin D and not vehicle (mixed-way ANOVA: Test X Treatment (F^(1,16)^=9.97, p<0.01), post-hoc: CPP test 2 (p<0.05)). *p<0.05, **p<0.05; n=9 per group).

### Retrieval of alcohol-related memories causes a time-dependent upregulation of *Arc* and *Zif268* but not *Bdnf* mRNA expression in the DH and mPFC

We next assessed whether alcohol memory retrieval alters the expression of the genes previously implicated in memory reconsolidation, namely activity-regulated cytoskeleton-associated protein (*Arc*)^5, 21, 36, 37^, transcription factor *Zif268* (also known as *Egr1*)^18, 20, 38^, and brain-derived neurotrophic factor (*Bdnf*)^39^, in the DH and mPFC, brain regions implicated in the reconsolidation of drug memories^5, 23, 24, 29, 30, 40^. To assess *Arc, Zif268*, and *bdnf* mRNA expression following alcohol memory retrieval, we first trained mice for alcohol-CPP (Figure 2A-B). Twenty-four hours later, mice were re-exposed to the alcohol-paired compartment (Retrieval group) or were handled (No Retrieval group). Brain tissues were collected at five different time points after memory retrieval, and target mRNAs levels were analyzed.

**Figure 2.**
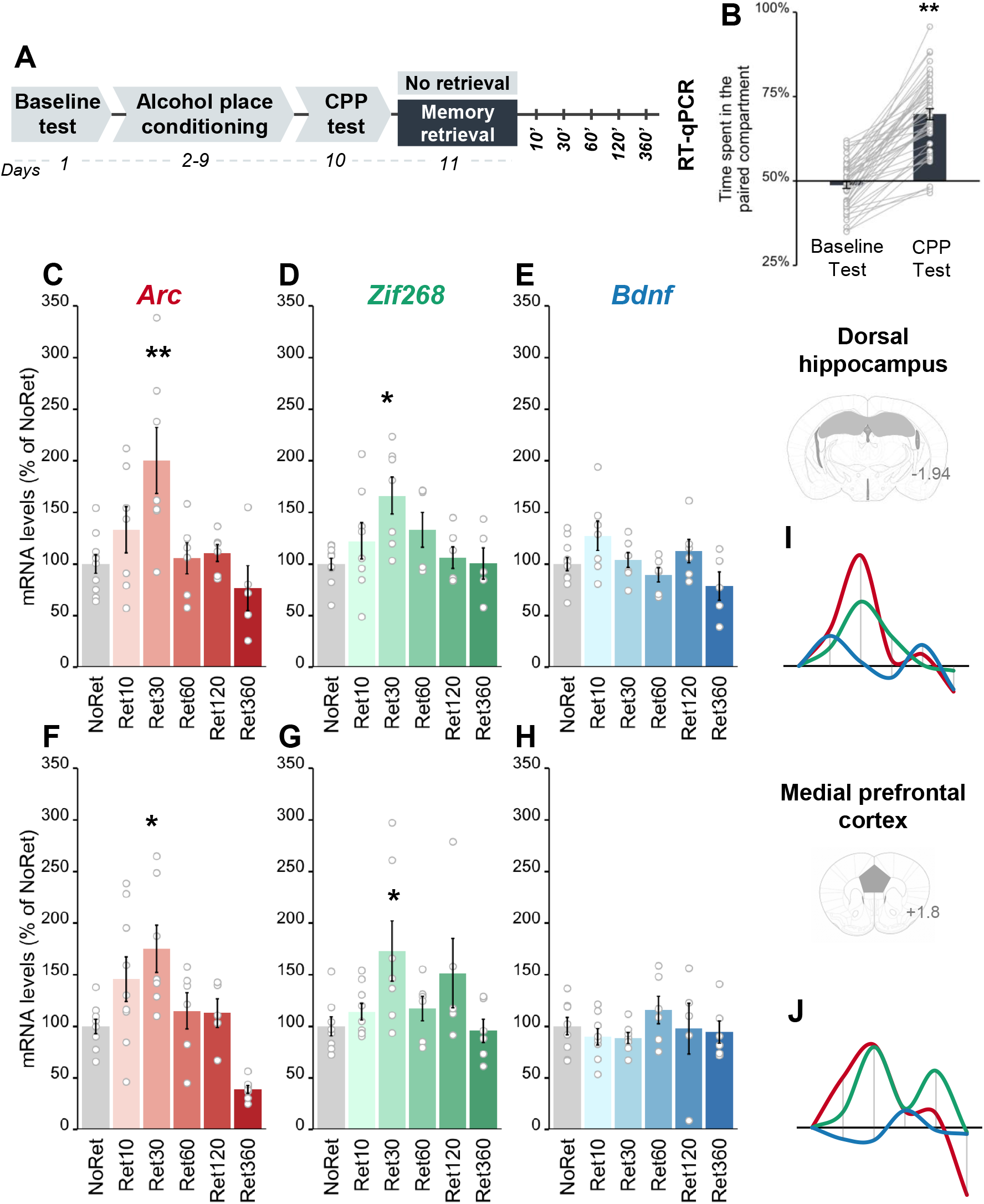
Alcohol memory retrieval triggers upregulation of *Arc* and *Zif268* but not *Bdnf* mRNA expression in the dorsal hippocampus and medial prefrontal cortex. **A**. Schematic illustration of the experimental design and timeline. **B**. Place preference scores, expressed as means ±S.E.M. of the percent of time spent in the alcohol-paired compartment (t^(47)^=13.82, p<0.0001); **C-H**. mRNA levels, normalized to *Gapdh*, of the percent of change from the control group (No Retrieval). qRT-PCR analysis revealed post-retrieval alterations in gene expression (one-way MANOVA; DH: Time (F^(15,86)^=2.42, p<0.01); mPFC: Time (F^(15,86)^=2.96, p<0.001): time-dependent upregulation of mRNA levels of *Arc* in the DH (NoRet vs Ret30’: p<0.01) (C) and mPFC (NoRet vs Ret30’: p<0.05) (F), of *Zif268* in the DH (NoRet vs Ret30’: p<0.05) (D) and the mPFC (NoRet vs Ret30’: p<0.05) (G), but not of *Bdnf* in the DH (E) or mPFC (H) (all p’s>0.05); **I-J**. Schematic representation of the time-dependent expression of *Arc* (red), *Zif268* (green), and *Bdnf* (blue) mRNA in the DH (**I**) and mPFC (**J**). Data are expressed as means ±S.E.M. *p<0.05; **p<0.01; n=9-6 per group.

Alcohol memory retrieval triggered rapid and transient upregulation in mRNA expression of *Arc* and *Zif268*, but not of *Bdnf* in the DH (Figures 2C-E and 2I). Specifically, *Arc* and *Zif268* mRNA levels peaked 30 min after alcohol memory retrieval, and returned to baseline levels within 60 min after memory retrieval, much like the No Retrieval group. In the mPFC, alcohol memory retrieval caused transient upregulation in *Arc* and *Zif268* but not *Bdnf* mRNA expression, similar to the expression pattern seen in the DH (Figures 2G-H and 2J). The increases in *Arc* and *Zif268* mRNA expression in the DH and mPFC were preceded by increased phosphorylation of the transcription factor cAMP response element-binding protein (CREB) (Figure S2), previously shown to regulate the expression of these genes^41^.

Together, the results show that the retrieval of alcohol-related memories induced a time-dependent upregulation in the expression of *Arc* and *Zif268* but not of *Bdnf* in the DH and mPFC, raising the possibility that altered expression of these genes may be involved in the reconsolidation of alcohol memories.

### The retrieval of alcohol-associated memories increases ARC protein levels in the DH

We previously showed that alcohol memory retrieval increased ARC protein levels in the amygdala and mPFC^5^. We now asked whether ARC protein levels were also increased in the DH, given the upregulation of *Arc* mRNA induced by alcohol memory retrieval (Figure 2C). Accordingly, mice were trained to express alcohol-CPP, as described above (Figures 3A-B). A day after the CPP test, alcohol memories were retrieved, and brain tissues were collected 60, 120 or 360 min later. We found that ARC protein levels in the DH increased 60 min after alcohol memory retrieval, returning to baseline levels within the next hour (Figures 3C and S3). These results thus suggest that alcohol memory retrieval increases *Arc* mRNA and ARC protein expression in the DH.

**Figure 3.**
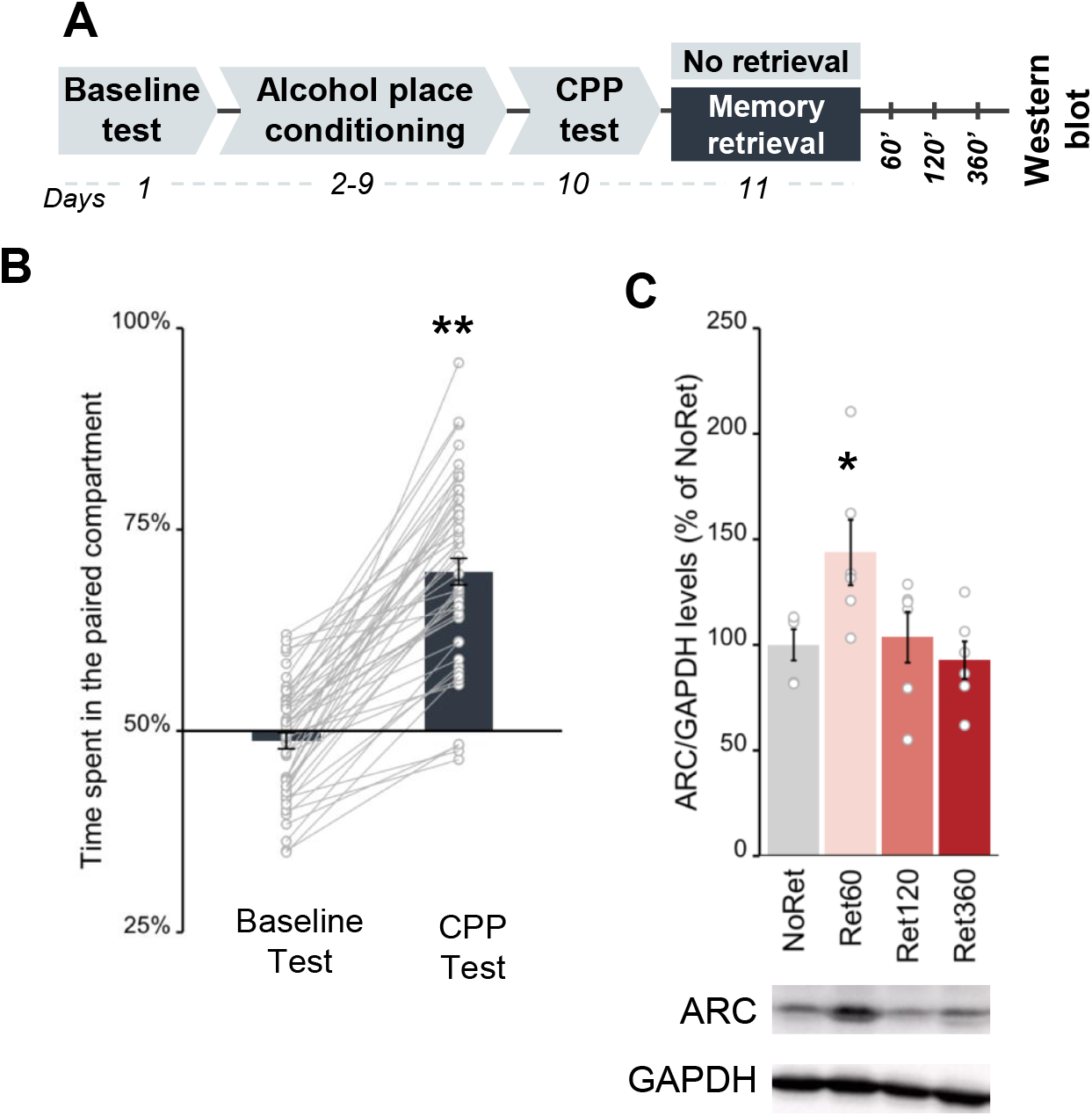
Alcohol memory retrieval induces upregulation of ARC protein expression in the DH. **A**. Schematic illustration of the experimental design and timeline. **B**. Place preference scores, expressed as means ±S.E.M. of the percent of time spent in the alcohol-paired compartment (t^(25)^=9.41, p<0.0001); **C**. ARC protein levels, normalized to GAPDH, expressed as means ±S.E.M. of the percent of change from the control group (No Retrieval). The levels of ARC protein in the DH were increased 60 min after alcohol memory retrieval (one-way ANOVA; Time (F^(3,19)^=3.94, p<0.05); post hoc: NoRet vs Ret60’ (p<0.05)). *p<0.05; **p<0.01; n=6-7 per group.

### Downregulation of ARC expression in the DH disrupts alcohol memory reconsolidation

If the increase of ARC expression in the DH following alcohol memory retrieval is essential for alcohol memory reconsolidation, then its downregulation following alcohol memory retrieval should disrupt such memory, resulting in the abolition of alcohol-CPP expression. To downregulate ARC levels in the DH during alcohol memory reconsolidation, we used antisense oligodeoxynucleotides (AS-ODN) directed against *Arc* mRNA^42^. Knockdown of ARC in brain regions related to memory consolidation and reconsolidation using *Arc* AS-ODN was previously shown to disrupt the consolidation of aversive and appetitive memories^36, 37, 42, 43^ and the reconsolidation of fear memories^36, 37, 43^, as well as to impair morphine-associated memory reconsolidation^21^.

To test whether ARC downregulation disrupts alcohol memory reconsolidation and abolishes alcohol seeking, we trained mice to show alcohol-CPP (Figure 4B, see Figure S5B for individuals’ data). A day after CPP test 1, the mice received an intra-hippocampal infusion of *Arc* AS-ODN or control scrambled (SCR)-ODN. Since the AS-ODN downregulated ARC protein levels 5 h after infusion^43^ (Figure S4, AS-ODN validation), alcohol memory was retrieved 4 h after infusion, allowing the downregulation to occur an hour after retrieval, at around the peak of increase in ARC protein levels induced by alcohol memory retrieval (Figure 3C). When place preference was tested the next day, we found that alcohol-CPP was abolished in mice that had received *Arc* AS-ODN, whereas SCR ODN-treated mice still presented CPP (Figure 4B). Our findings thus suggest that intra-hippocampal infusion of *Arc* AS-ODN disrupted the reconsolidation of alcohol memories by preventing the increases of ARC protein levels caused by memory retrieval.

**Figure 4.**
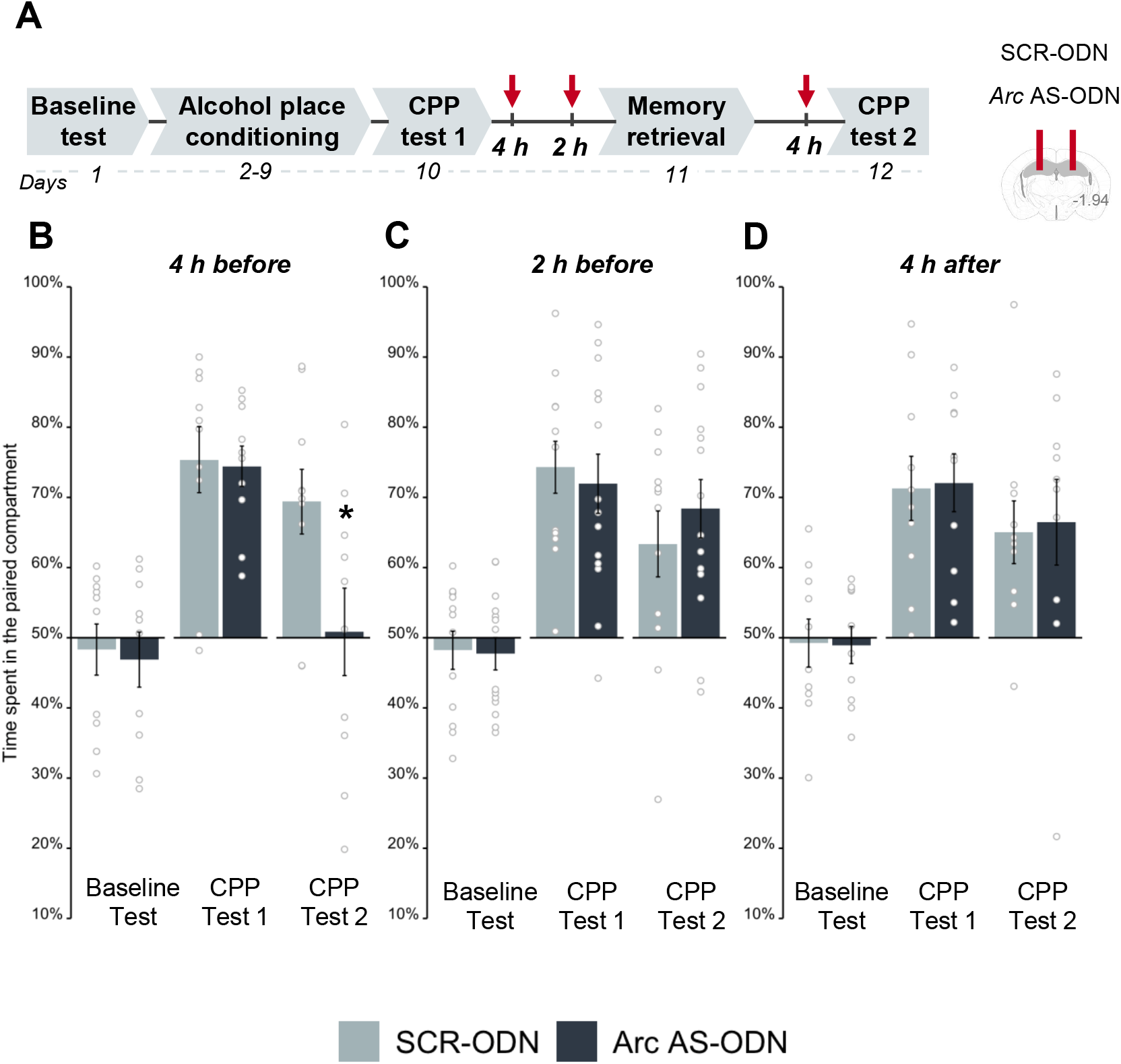
Downregulation of ARC protein expression in the dorsal hippocampus shortly after alcohol memory retrieval disrupts the expression of alcohol-conditioned place preference (CPP). **A**. Schematic illustration of the experimental design and timeline. Antisense oligodeoxynucleotides directed against *Arc* mRNA (*Arc* AS-ODN) or non-specific scrambled oligodeoxynucleotides (SCR-ODN) were infused into the dorsal hippocampus (DH) of mice at the indicated time points. **B-D**. Place preference scores, expressed as means ±S.E.M. of the percent of time spent in the alcohol-paired compartment. Infusion of *Arc* AS-ODN disrupted the expression of alcohol-CPP when infused 4 h (mixed-model ANOVA; Test (F^(1,18)^=38.04, p<0.001), and Test X Treatment (F^(1,18)^=13.48, p<0.01); post hoc: CPP test 2 (p<0.05)) **(B)** but not 2 h before memory retrieval (all p’s>0.05) **(C)** or 4 h after memory retrieval (all p’s>0.05) **(D)**. *p<0.05; n=10-12 per group.

To further test whether this memory disruption was due to the blockade of the post-retrieval ARC induction, we chose a second time point for *Arc* AS-ODN infusion, one still within the “reconsolidation window”, yet when ARC protein expression levels had returned to baseline (as indicated in Figure 3C). Thus, we infused *Arc* AS-ODN or control SCR-ODN into the DH 2 h before memory retrieval, which is expected to downregulate ARC levels 5 h later, i.e., 3 h after memory retrieval (Figure S4B). In a place preference test conducted a day later, we found that both groups persisted in showing alcohol-CPP (Figure 4C). These findings indicate that downregulation of ARC levels past its retrieval-dependent induction does not interfere with the ongoing reconsolidation of alcohol memories.

We further assumed that downregulation of ARC outside the “reconsolidation window”, i.e., more than 5-6 h after memory retrieval ^9, 10^, would not affect subsequent memory expression. To test this hypothesis, we infused *Arc* AS-ODN or control SCR-ODN into the DH 4 h after memory retrieval, which is expected to affect ARC protein expression 9 h after memory retrieval (i.e., 5 h later; Figure S4). Testing a day later revealed that both groups demonstrated strong preferences for the alcohol-paired compartment. These findings indicate that downregulation of ARC protein expression several hours after memory retrieval (i.e., outside the reconsolidation window) does not affect the memories underlying the expression of alcohol-CPP.

Together, these findings suggest that the hippocampal upregulation of ARC protein expression observed shortly after alcohol memory retrieval is required for reconsolidating alcohol memories, as inhibition of these retrieval-induced increases of ARC protein levels led to the loss of alcohol seeking, likely by disrupting the memory reconsolidation process.

### Upon retrieval, appetitive alcohol-and non-alcohol-associated memories share similar *Arc* and *Zif268* transcriptional dynamics

Our findings impllicating *Arc* and *Zif268* expression in the reconsolidation of alcohol memories are in line with previous studies showing these IEGs are implicated in the reconsolidation of different types of memory^18, 21, 36, 37, 43^. We assumed that the transcription and translation of these IEGs are not specific for alcohola, and rather may play a part in the common basic mechanisms for the processing of reactivated memories, including appetitive memories^15^. To further explore this possibility, we tested whether the retrieval of non-alcohol, sucrose-associated memories via a similar CPP protocol would alter *Arc* and/or *Zif268* mRNA expression, as it did with alcohol-related memories. For this, we first trained mice in a sucrose-CPP procedure similar to the alcohol-CPP procedure used above, pairing one compartment of the CPP apparatus with voluntary consumption of sucrose pellets (Figure 5A, Experimental design). After four pairings, mice showed strong preference for the sucrose-associated compartment (Figure 5B). Next, the sucrose-associated memory was retrieved by re-exposure to the sucrose-paired compartment. Brain tissues from the Retrieval and No retrieval control groups were collected 10, 30 or 60 min after memory retrieval. *Arc* and *Zif268* mRNA levels in the DH and mPFC were then assessed.

**Figure 5.**
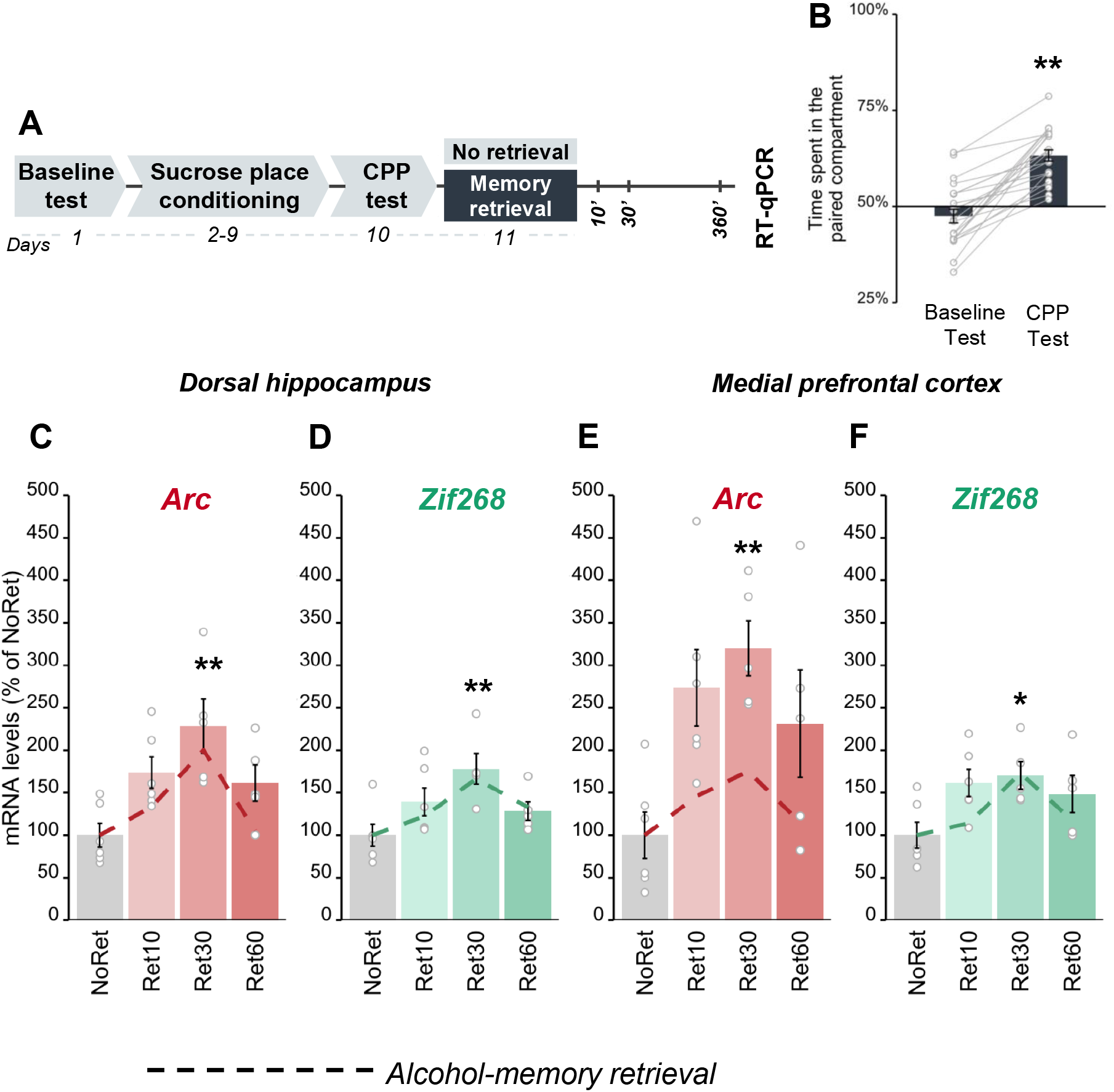
Sucrose memory retrieval triggers upregulation of *Arc* and *Zif268* in the dorsal hippocampus and medial prefrontal cortex. **A**. Schematic illustration of the experimental design and timeline. **B**. Place preference scores, expressed as means ±S.E.M. of the percent of time spent in the sucrose-paired compartment (t^(21)^ = 8.45,p<0.0001); **C-F**. mRNA levels, normalized to *Gapdh*, expressed as means ±S.E.M. of the percent of change, as compared with the control group (No Retrieval). qRT-PCR analysis revealed post-retrieval alterations in gene expression (one-way MANOVA; DH: Time (F^(6,34)^=2.82, p<0.05); mPFC: Time (F^(6,34)^=2.13, p=0.07)). *Arc* mRNA levels were transiently increased in the DH (NoRet vs Ret30’: p<0.01) (C) and mPFC (NoRet vs Ret30’: p<0.01) (E), while *Zif268* expression was increased in the DH (NoRet vs Ret30’: p<0.01) and the mPFC (NoRet vs Ret30’: p<0.05) (F). mRNA expression levels after alcohol memory retrieval are shown as dashed lines. *p<0.05; **p<0.01; n=9-6 per group.

We found that sucrose memory retrieval caused rapid and transient upregulation of *Arc* and *Zif268* mRNA expression in both the DH and mPFC (Figure 5C-F). As depicted by the dashed lines in Figure 5C-F, the patterns of mRNA expression upregulation in both brain regions, induced by the retrieval of sucrose memories, resembled the upregulation of these genes observed upon alcohol memory retrieval. These findings thus suggest that rapid and transient upregulation in *Arc* and *Zif268* mRNA expression in the DH and mPFC is associated with the retrieval of both alcohol-and sucrose-associated memories.

### Characterization of alcohol-specific transcriptional dynamics for memory retrieval: A transcriptomic analysis

Given our finding that *Arc* and *ZIf268* transcription were altered upon retrieving both alcohol-and non-alcohol-associated memories, we next sought to identify the transcriptomic signature specific for alcohol memory retrieval by performing RNA-seq analysis of the DH and mPFC (Figure 6A, Experimental design). To this end, following alcohol-CPP training (Figure 6B), alcohol-associated memories were retrieved (with a No retrieval control group). The DH and mPFC were collected 30 min later and processed for RNA-seq analysis (Figure 6C-D).

**Figure 6.**
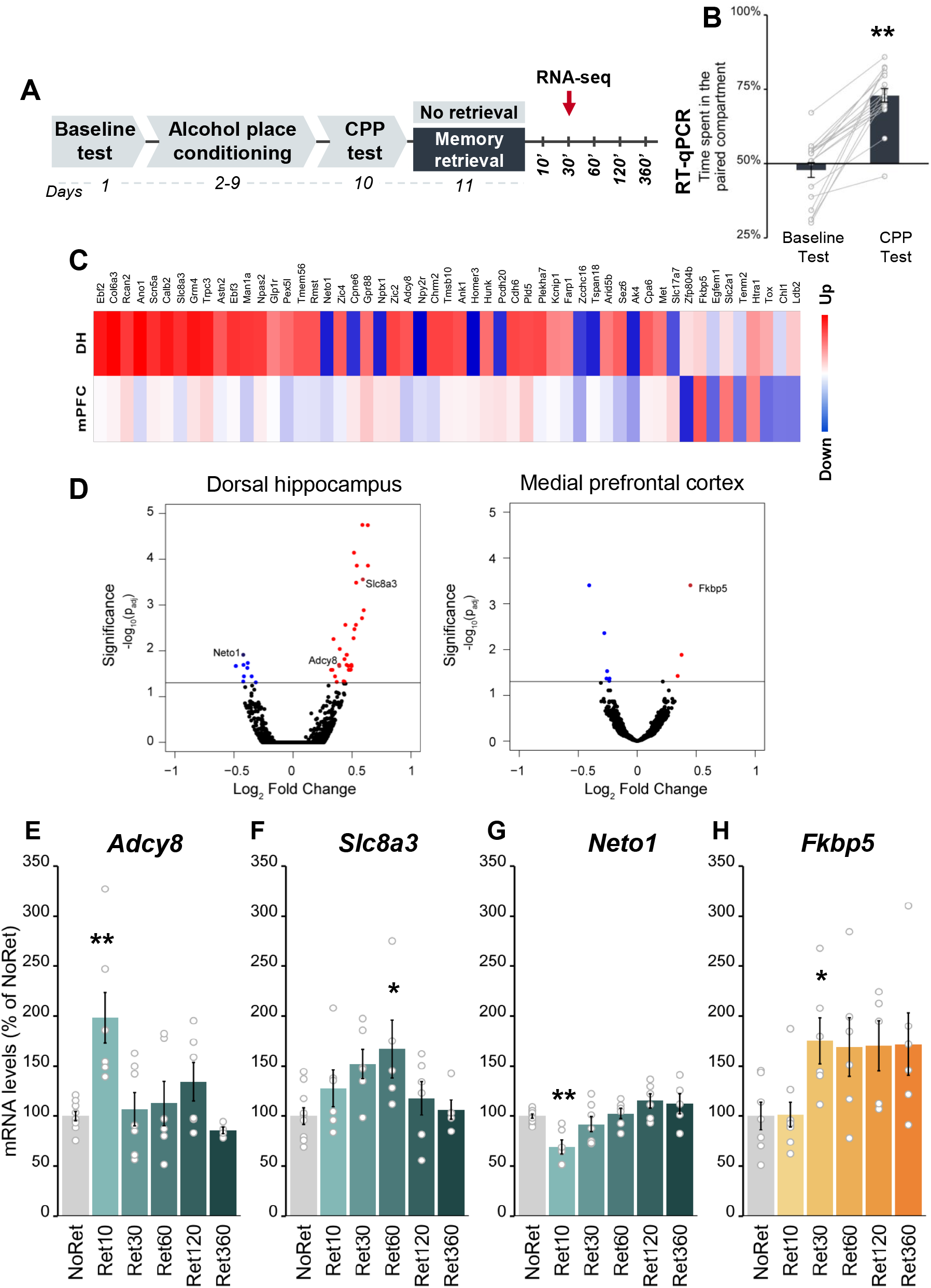
Alcohol memory retrieval alters transcriptomic dynamics in the dorsal hippocampus and medial prefrontal cortex. **A**. Schematic illustration of the experimental design and timeline. **B**. Place preference scores, expressed as means +/-S.E.M. of the percent of time spent in the alcohol-paired compartment (t^(17)^=8.36, p<0.0001); **C**. A heat map generated by hierarchical analysis of genes identified using DESeq2 shows significant changes in expression in the mPFC and/or DH following alcohol memory retrieval, as compared to the control No Retrieval group, with a significance cutoff-adjusted p-value (p^(adj)^< 0.05). Red=up-regulated genes; blue=down-regulated genes. **D**. A volcano plot provides an overview of the genes detected by RNA-sequencing. Log2-fold changes are plotted on the x-axis, and the negative log10 (p-value) is plotted on the y-axis. Differentially expressed genes appear above the line that indicates the significance threshold. Red=up-regulated genes; blue=down-regulated genes. **E-H**. mRNA levels, normalized to *Gapdh*, expressed as means ±S.E.M. of the percent of change from the control group (No Retrieval). qRT-PCR analysis revealed wave-like alternations in the levels of selected genes, detected via RNA-sequencing. In the DH: *Adcy8* (one-way ANOVA; Time (F^(5,33)^=5.08, p<0.01); post-hoc: NoRet vs Ret10 (p<0.01)) (**E**), *Slc8a3* (Time (F^(5,33)^=2.78, p<0.05); post-hoc: NoRet vs Ret60 (p<0.05)) (**F**), *Neto1* (Time (F^(5,33)^=6.27, p<0.01); post-hoc: NoRet vs Ret10 (p<0.01)) (**G**). In the mPFC: *Fkbp5* (Time (F^(5,33)^=3.07, p<0.05); post-hoc: NoRet vs Ret30 (p<0.05)) (**H**). *p<0.05, **p<0.01; n=10-6.

Using DESeq2^44^, we identified a set of 44 genes whose levels of expression were significantly altered in the DH, with 34 genes being upregulated, and 10 genes being downregulated (Figure 6C-D and Table S2). We further found that in the mPFC, the expression of 9 genes was significantly altered (3 were upregulated and 6 were downregulated); none of these genes overlapped with those DH genes showing altered expression (Figure 6C). The differential expression of selected genes detected by RNA-seq was further confirmed by quantitative reverse transcriptase-polymerase chain reaction (qRT-PCR) analysis of brain samples collected at five different time points following memory retrieval (Figure 2A-B). Memory retrieval led to downregulation of the mRNA expression of *Adcy8* (encoding adenylate cyclase 8) and *Slc8a3* (encoding solute carrier family 8 (sodium/calcium exchanger), member 3), and to downregulation of *Neto1* (encoding neuropilin (NRP) and tolloid (TLL)-like 1) expression in the DH (Figure 6E-G), as well as to upregulation of *Fkbp5* expression (encoding FK506 binding protein 5). The RNA-seq analysis also revealed mild upregulation of *Arc* and *Zif268 mRNA* expression in the DH and mPFC (Table S2), consistent with the earlier RT-qPCR results.

We next tested whether the expression of specific genes we found to be altered upon retrieval of alcohol memories were affected by the retrieval of non-alcohol, sucrose-associated memories, in a manner similar to the common upregulation of *Arc* and *Zif268* mRNA expression (Figure 7A-B). As shown in Figure 7C-F, *Adcy8, Slc8a3, Neto1* and *Fkbp5* mRNA expression was not affected by sucrose memory retrieval, although a trend towards increased expression of *Neto1* mRNA was noted after 10 min of sucrose memory retrieval, i.e., in the opposite direction to the decreased mRNA expression induced by alcohol memory retrieval.

**Figure 7.**
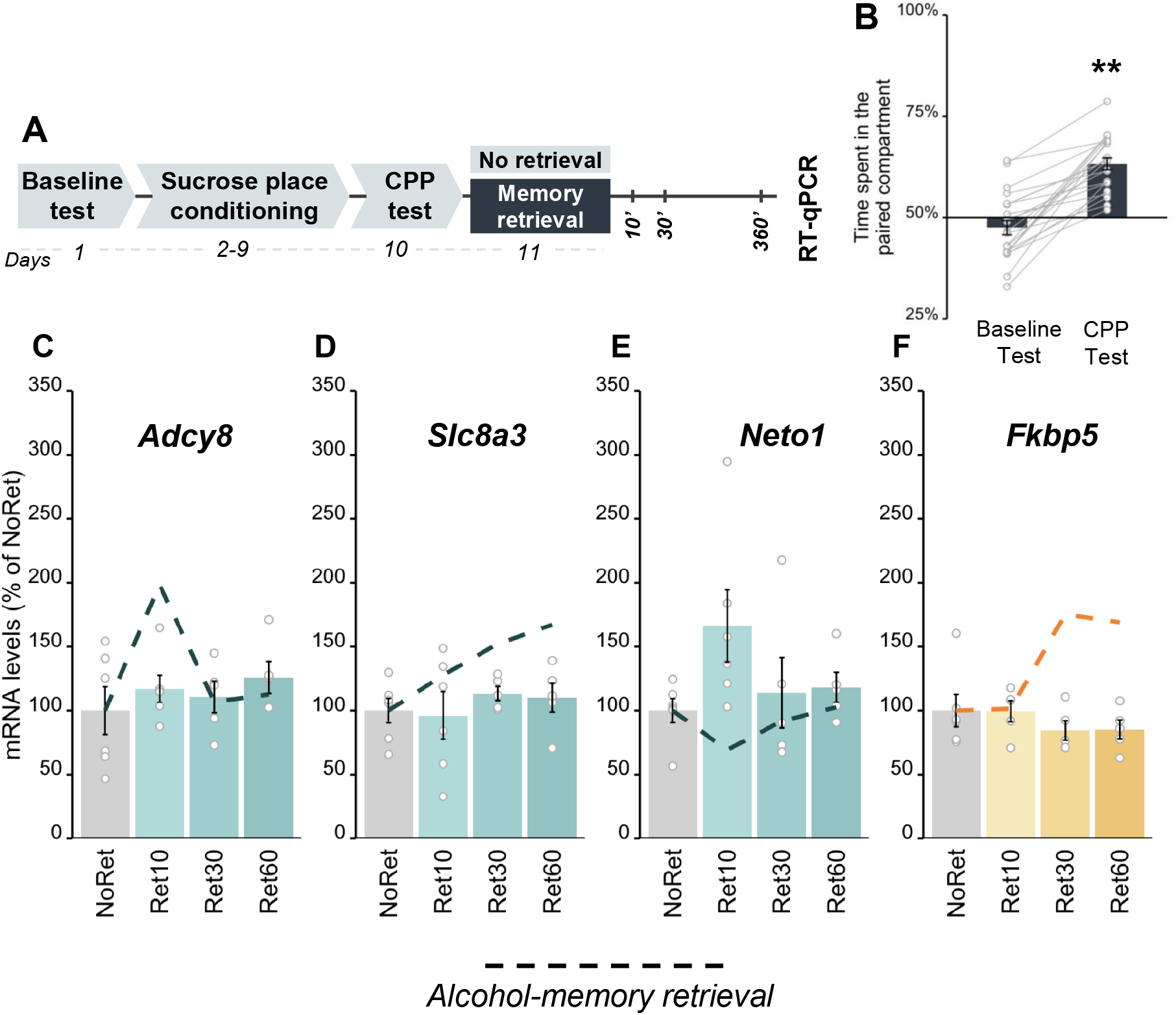
Sucrose memory retrieval does not alter the expression of *Adcy8, Slc8a3*, or *Neto1* in the dorsal hippocampus or of *Fkbp5* in the medial prefrontal cortex. A. Schematic illustration of the experimental design and timeline. **B**. Place preference scores, expressed as means ±S.E.M. of the percent of time spent in the sucrose-paired compartment (t^(21)^ = 8.45,p<0.0001); **C-F**. mRNA levels, normalized to *Gapdh*, expressed as means ±S.E.M. of the percent of change from the control group (No Retrieval). qRT-PCR analysis did not reveal differences in the mRNA levels of *Adcy8, Slc8a3, or Neto1* (**A-C**), and *Fkbp5* (**D**) in the DH and mPFC, respectively, between the Retrieval and No retrieval groups (one-way ANOVA; all p’s>0.05). mRNA expression levels after alcohol memory retrieval (presented in Figure 6E-H)) are shown as dashed lines; **p<0.01; n=9-6 per group.

To summarize, these experiments identified a unique transcriptional dynamics triggered by alcohol memory retrieval, which is not common for memories of a natural reward, namely, sucrose-associated memories.

## Discussion

We show here that the reconsolidation of alcohol-associated memories requires *de novo* gene transcription, and that alcohol seeking can be disrupted by inhibiting transcription following alcohol memory retrieval. Importantly, our findings suggest that while the altered expression of some genes is likely a common mechanism for the reconsolidation of several types of memories, the processing of alcohol-related memories is also characterized by a unique transcriptional profile.

Specifically, we found that the retrieval of either alcohol memories or of memories associated with a natural reward (sucrose) triggered similar increases in mRNA expression of the IEGs *Arc* and *Zif268* in the DH and mPFC. In contrast, RNA-seq analysis revealed a subset of genes (*Adcy8, Slc8a3*, and *Neto1* in the DH, and *Fkbp5* in the mPFC) of which expression was altered selectively by the retrieval of alcohol, but not by sucrose reward memories, raising the intriguing possibility that alcohol-associated memories that trigger relapse have unique molecular mechanisms that could be targeted to disrupt them selectively.

We show that the downregulation of ARC shortly after alcohol memory retrieval abolishes alcohol seeking (the expression of alcohol CPP), indicating a critical role for hippocampal ARC expression in the reconsolidation of alcohol memories. *Arc* is a CREB-regulated IEG, rapidly induced by neuronal activity, and known to regulate synaptic plasticity and mediate memory formation^42^. We previously showed that inhibition of the translational machinery that controls ARC protein synthesis disrupted alcohol memory reconsolidation and prevented relapse to alcohol seeking and drinking in a rat self-administration paradigm^5^. Indeed, *Arc* has been described as a key player in the reconsolidation of drug^5, 12, 21^-and fear^36, 37^-associated memories.

We found that *Arc* mRNA expression peaked 30 min after alcohol memory retrieval, and returned to baseline levels within the next 30 min, followed by a transient increase in ARC protein levels, peaking 1 h after memory retrieval. These transient ARC dynamics imply that alcohol memory retrieval induces rapid mRNA^45^ and protein^46^ degradation. Accordingly, we found that alcohol memory reconsolidation was disrupted when we knocked-down the expression of ARC 1 h but not 3 or 9 h after memory retrieval. It is accepted that the “reconsolidation window” lasts ∼5-6 h, as manipulations conducted 5-6 h after memory retrieval failed to affect targeted behaviors^7, 10, 11^. Our results, therefore, suggest that the reconsolidation window might be in fact narrower than 5 hours, at least when ARC protein expression up-regulation is required for memory re-stabilization.

While we localized the causal role of ARC in alcohol memory reconsolidation to the DH, a brain region previously implicated in the reconsolidation of contextual drug memories^29, 30, 38, 40^, *Arc* and *Zif268* mRNA expression was also upregulated in the mPFC. Both the prelimbic and infralimbic prefrontal subregions have been implicated in drug memory reconsolidation^23, 24^, suggesting the mPFC to be a candidate brain region that regulates alcohol seeking via reconsolidation mechanisms. Our current finding is consistent with our previous observation of upregulated ARC protein levels in this brain region following alcohol memory retrieval in an operant alcohol self-administration procedure^5^.

In addition to *Arc*, we found that the mRNA levels of *Zif268*, but not *Bdnf*, were increased in the DH and mPFC by alcohol memory retrieval. Increased *Zif268* mRNA expression has been previously implicated in the reconsolidation of fear and drug memories^18, 20, 38, 47^, raising the possibility that this transcription factor also plays a role in the reconsolidation of alcohol memories. Whereas the role of *Bdnf* in memory reconsolidation remains controversial^18, 39^, *Bdnf* induction is known to be crucial for memory acquisition and extinction learning^18,48^. Thus, the lack of change in *Bdnf* expression in our study could suggest that our memory retrieval procedure did not initiate extinction learning in parallel with memory retrieval. Moreover, we recently reported that although the mRNA levels of *Bdnf* in the DH are not increased by alcohol memory retrieval *per se*, the expression of the growth factor were elevated when the retrieval is followed by aversive counterconditioning that prevented relapse^34^, suggesting a complex role of *Bdnf* in alcohol memory dynamics.

Our findings indicate that the increases in the IEGs expression are not unique to the reconsolidation of alcohol-related memories. Indeed, ARC and ZIF268 were previously implicated in the reconsolidation of several types of memories, including fear memories^18, 36, 37, 48^, recognition memories^20^, and memories associated with different drugs of abuse^19, 21, 47^. However, there is also evidence that some of the mechanisms underlying alcohol seeking may differ from those controlling natural reward seeking^5, 49-51^. Furthermore, there is evidence that memories for different rewards (and for different drugs, in particular) are differentially processed^52-54^. Our RNA-seq analysis findings indeed revealed the unique transcriptomic signature induced by alcohol memory retrieval. Specifically, we found 44 genes in the DH and a different set of 9 genes in the mPFC, which showed significant changes following alcohol memory retrieval. Following further data validation, we focused on one gene in the mPFC (*Fkbp5*) and three genes in the DH (*Adcy8, Scl8a3, and Neto1*) and found that their expression was not affected by the retrieval of sucrose-associated memories. This suggests that unlike *Arc* and *Zif268*, these 4 specific genes likely do not play general roles in the processing of reward memories, yet may rather play selective roles in the processing of alcohol memories.

Interestingly, these four genes were previously implicated in alcohol use disorder and in learning and memory. For example, *Fkbp5* encodes the FK506 Binding Protein 5, a regulator of the stress-neuroendocrine system^55^. *FKBP5* variants modulate the severity of alcohol withdrawal syndrome^56^, and predict the propensity of heavy drinking in humans^57-59^. Deletion of *Fkbp5* was shown to increase alcohol withdrawal severity^56^ and alcohol drinking^58^ in mice, whereas pharmacological inhibition of the protein reduced moderate alcohol consumption, and reinstatement of CPP^60^. The levels of *Adcy8* mRNA, encoding adenylate cyclase 8 (AC8) that catalyzes cAMP formation in response to calcium influx and recently marked as a possible regulator of alcohol intake^51, 61^, were decreased in blood cells from long-term abstinent alcoholics^62^. Deletion of *Adcy8* was previously shown to reduce alcohol drinking and increase sensitivity to the sedative effects of alcohol in mice^63^. In addition, chronic alcohol exposure in rats upregulated the brain expression of solute carrier family 8 (sodium/calcium exchanger), member 3 (NCX3), the protein encoded by *Scl8a3*^64, 65^. Alcohol consumption in rats was also associated with reduced brainstem expression of *Neto1*^66^, encoding neuropilin tolloid-like 1 (NRP1), a component of the NMDA-receptor complex^67^. This protein is involved in synaptic reorganization and transmission in the hippocampus^68^, and is required for spatial learning and memory^67^. Thus, our findings show that the mere retrieval of alcohol memories, even without any pharmacological effects of alcohol, affect the expression of these genes. However, their specific role in alcohol memory reconsolidation and relapse remains to be tested.

In summary, our findings suggest that alcohol memory retrieval induces two parallel transcription programs. One program, conveyed via common molecular mechanisms of learning and memory, including the IEGs *Arc* and *Zif268*, is engaged in the reconsolidation of memories, in general. The other transcription program, launched by alcohol memory retrieval, is controlled by genes that specifically promote alcohol-related behaviors. This dual-processing model for alcohol memories raise the possibility that memory reconsolidation for different memories may have similar dual-processing molecular mechanisms, with some components shared by multiple memories, and others unique to each memory type. As such, this hypothetic model suggests that it may be possible to aim at reward-specific molecular targets to treat disorders related to pathogenic memories, such as addiction, rather than disrupt major molecular mechanisms essential for many functions beyond alcohol memory reconsolidation.

## Materials and methods

### Animals

Male and female C57BL/6 mice (25-30 g), housed 3-4/cage were bred at the Tel-Aviv University Animal Facility (Israel), and kept under a 12 h light-dark cycle (lights on at 7 a.m.), with food and water available *ad libitum*. In experiments involving sucrose place-conditioning, access to food was restricted to 15-16 h a day: the food tray was emptied 6 h prior to the beginning of training and 1 h after the lights were turned on, and refilled 1 h after the end of the manipulations; water was supplied *ad libitum*. A food restriction schedule was launched 3 days before sucrose training and lasted until the last day of the experiment (20 days in total). Mice were weighed twice a week to control for weight loss. All experimental protocols were approved by and conformed to the guidelines of the Institutional Animal Care and Use Committee of Tel Aviv University, and to NIH guidelines of the (animal welfare assurance number A5010-01). All efforts were made to minimize the number of animals used.

### Drugs and reagents

Ethanol absolute (Gadot, Israel) was diluted with tap water to a 20% (v/v) solution for voluntary intake or in sterile saline solution (0.9 M NaCl) for systemic injections. Actinomycin D was purchased from Sigma-Aldrich (Rehovot, Israel) and dissolved in DMSO (Sigma-Aldrich, Rehovot, Israel)). Isoflurane was obtained from Piramal Critical Care (Bethlehem, PA, USA). Fast SYBR Green Master Mix, TRIzol reagent and RevertAid kit were supplied by Thermo-Fisher Scientific (Waltham, MA, USA). DNA oligonucleotides (RT-qPCR primers) were obtained from Sigma-Aldrich (Rehovot, Israel). Mouse monoclonal antibodies against ARC (sc-17839) and GAPDH (sc-32233) were purchased from Santa Cruz Biotechnology (Santa Cruz, CA, USA). Nitrocellulose membranes were purchased from Millipore (Billerica, MA, USA). Protease and Phosphatase Inhibitor Cocktail (100x) and enhanced chemiluminescent (ECL)-horseradish peroxidase (HRP)HRP substrate were purchased from Thermo Fisher Scientific (Kiryat Shmona, Israel).

### Apparatus

#### Place conditioning

All place conditioning experiments were performed in open ceiling Plexiglas boxes (30 × 30 × 20 cm) divided into two equal-sized compartments by a sliding door. The compartments differed from each other in terms of wall pattern (horizontal vs. vertical black and white stripes, 1 cm-wide) and floor surface (white textured plastic with bulging circles vs. bulging stripes). The horizontal striped pattern on the walls was always matched with the bulging circles on the floor, whereas the vertical stripes were linked with the bulging stripes. Each Plexiglas box was placed in a sound-attenuating chamber equipped with a LED light stripe on the walls and a ceiling camera that registered mouse behavior. Data were recorded with an Ethovision XT 11.5 video tracking system (Noldus, Wageningen, Netherlands).

### Behavioral procedures

#### Place conditioning

##### Alcohol-conditioned place preference (CPP)

All mice were habituated to daily i.p. saline injections 5 days prior to the beginning of the procedure.

##### Baseline test (day 1)

On the first day, the sliding door was retracted and mice were allowed to freely explore the entire apparatus for 30 min. Animals that spent >75% of time in either compartment were excluded from the study. This allowed the use of an unbiased design, in which the two compartments were equally preferred before conditioning, as indicated by the group average (unbiased apparatus), and pseudo-randomly assigned to the experimental conditions (unbiased assignment procedure) ^31^. Each place-conditioning chamber was assigned to a single sex.

##### Alcohol place conditioning (days 2–9)

Training started 24 h after the baseline test with one session administered per day over 8 days, with the sliding door closed. On days 3, 5, 7, and 9, the mice received alcohol (1.8 g/kg, 20% v/v; i.p.) and immediately confined to the paired compartment for 5 min. This dose and conditioning duration were previously shown to produce alcohol-induced CPP ^32, 69^. On the alternate days (i.e., days 2, 4, 6, and 8), the mice received saline solution and were confined to the unpaired compartment for the same duration as on the alcohol-conditioning day. Paired compartments were counterbalanced.

##### Place preference test 1 (day 10)

Place preference testing was as described for the baseline test, and served to index alcohol-CPP ^31^. Preference was defined as an increase in the percent of time spent in the alcohol-paired compartment during place preference test 1, as compared to the baseline test.

##### Memory retrieval (day 11)

Prior to this stage, the mice were assigned to different experimental conditions (matched for CPP scores and sex). During a memory retrieval session, the mice were confined to the alcohol-paired compartment for 3 min, and then returned to their home cages. Control mice were handled briefly.

##### Place preference test 2 (day 12)

The mice were subjected to a place preference test identical to place preference test 1.

##### Sucrose CPP

##### Sucrose pre-training (days 1-6)

On days 1-3, the mice were habituated to collect 5-6 sucrose pellets (45 mg, Dustless Precision Pellets, Bio-Serv, Frenchtown, NJ, USA) from a plastic plate inserted into the home cage. The plates were removed from the home cage when emptied. On days 4-6 (habituation for food in the apparatus), mice were brought to the experimental room, with the CPP boxes having been stripped of their wall patterns and textured floor covers. The sliding doors were fully retracted, such that the mice were allowed to freely explore the boxes. A min later, three sucrose pellets were scattered on the floor. Mice were removed from the boxes shortly after collecting all the pellets, or 30 min after the beginning of the session. Mice that collected fewer than six pellets (out of a total of nine pellets over the 3 days) during sucrose training were excluded from the experiment (2 mice out of 32). Following the pretraining days, the wall patterns and textured floor covers of the CPP apparatus were restored. The sliding door was inserted when indicated.

##### Baseline test (day 7)

On the seventh day, the sliding door was retracted and the mice were allowed to explore the entire apparatus for 30 min. An unbiased apparatus/unbiased assignment approach, as described in the alcohol-CPP procedure, was also adopted here.

##### Sucrose place-conditioning (days 8-15)

Training started 24 h after the baseline test with one session per day over 8 days, with the sliding door closed. On days 9, 11, 13, and 15, mice were placed in the paired compartment, and a min later, three sucrose pellets were scattered on the floor. Mice were returned to their home cages 15 min later. On the alternate days (i.e., days 8, 10, 12, and 14), mice were placed in the unpaired compartment for 15 min with no interference. Paired compartments were counterbalanced.

##### Place preference test 1 (day 16)

Place preference testing was as described for the baseline test, and served to index sucrose-CPP. Preference was defined as an increase in the percent of time spent in the sucrose-paired compartment during place preference test 1, as compared to the baseline test.

##### Memory retrieval (days 17)

The sucrose memory retrieval procedure was identical to the alcohol memory retrieval procedure (see above).

### Oligodeoxynucleotide (ODN) design, preparation and validation

*Arc* antisense ODN (AS-ODN) and scrambled ODN (SCR-ODN) (Sigma-Aldrich, Rehovot, Israel) design followed the guidelines described previously^42^. The *Arc* AS-ODN encoded an antisense sequence for the *Arc*/*Arg3*.*1* mRNA sequence that overlaps the translation start site, and was previously shown to knockdown *Arc* mRNA translation in mice^70^. In addition, it was reported to inhibit ARC protein expression in the hippocampus without affecting the translation of other genes^42, 70^ and to impair the reconsolidation of fear memories^36^. In a pilot study, we confirmed the ability of *Arc* AS-ODN to reduce ARC protein expression in the DH, as compared to a SCR-ODN control (∼40% reduction) (Figure S4). The SCR-ODN, containing the same nucleotides as the *Arc* AS-ODN but assembled in a random sequence did not show significant homology to sequences in the GenBank database and served as a control. Both ODNs contained phosphorothioate linkages at the bases of both the 5′ and 3′ ends and phosphodiester internal bonds, given how this nucleotide design was reported to be more stable than a unmodified phosphodiester ODN *in vivo* and less toxic than a fully phosphorothioated ODN ^42^. The following sequences were used (* denotes a phosphorothioate linkage): 5′-G*T*C*CAGCTCCATCTGGT*C*G*T-3′ (*Arc* AS-ODN) and 5′-C*G*T*GCACCTCTCGCAGG*T*T*T-3′ (SCR-ODN).

*Arc AS-ODN functional validation*: *Arc* AS-ODN (2 nmol/µl, 0.5 µl, 0.25 µl/min) were infused into the DH in one hemisphere, and SCR-ODN (2 nmol/µl, 0.5 µl, 0.25 µl/min) was infused into the other hemisphere (the sides were counterbalanced). Two or four hours later, treated mice explored an unfamiliar context (CPP compartment, see Apparatus description, above) for 5 min to induce novelty-dependent ARC expression. Mice were euthanized an hour later (i.e., 3 or 5 h after ODN infusion), brain tissues were collected, and ARC protein levels were assessed by Western blot analysis (Figure S4).

### Surgery and intra-hippocampal microinfusion

Surgery and microinfusions were conducted as described previously^32, 71^.

#### Surgery

Stereotaxic surgeries were conducted under isoflurane anesthesia. Mice were placed in a stereotaxic frame (RWD Life Science, Shenzhen, China) and bilateral guide cannulae (C235G-2.6, 26G; Plastics One Inc., Roanoke, VA, USA) were aimed at the DH at the following coordinates (−2 mm posterior to bregma, ±1.3 mm mediolateral, -1.45 mm ventral to the skull surface). Cannulae were secured with dental acrylic. Matching dummy cannulae (Plastics One Inc., Roanoke, VA, USA) were inserted into the guide cannulae and topped with dust cups to keep the injector site covered and clear of debris. The mice were allowed to recover for 7-10 d prior to alcohol-CPP training.

#### Intra-hippocampal infusions

Actinomycin D (4 µg/µl, 0.5 µl per side, 0.25 µl/min, in DMSO) ^18^ or an equivalent volume of vehicle was microinjected into the DH immediately after memory retrieval. Infusion of actinomycin D into the hippocampus at a similar concentration was shown to disrupt memory reconsolidation^18^. *Arc* AS-ODN or control SCR-ODN (2 nmol/µl, 0.5 µl/side, 0.25 µl/min; in PBS) was microinjected into the DH 4 h or 2 h before or 4 h after memory retrieval.

After removal of the dust cup and dummy cannulae, microinfusion was conducted over 2 min to awake gently restrained mice, using injection cannulae (33G; Plastics One Inc., Roanoke, VA, USA) extending 0.5 mm beyond the guide cannula tip. Injection cannulae were left in place for an additional 2 min. After infusion, dummy cannulae were inserted into the guide cannulae, secured with the dust cup, and the animals returned to their home cages. Cannulae locations were verified in 30 μm-thick coronal sections of paraformaldehyde-fixed tissue stained with cresyl violet.

### Western blot analysis

Western blot analysis was conducted as we previously described^5^. Briefly, brain tissues were rapidly dissected, snap-frozen with liquid nitrogen and stored at -80°C until use. Before Western blot analysis, samples were homogenized in buffer containing (in mM): 25 Tris-HCl, pH 7.6, 150 NaCl, 1 EDTA, 1% (v/v) NP-40, 0.5% (w/v) sodium deoxycholate, 0.1% (w/v) sodium dodecyl sulfate (SDS) and protease and phosphatase inhibitors. Protein concentrations were determined using a BCA assay, and equal amounts of each sample (40 μg) was denatured with Laemmli buffer, boiled for 10 min, resolved by 10% SDS–polyacrylamide gel electrophoresis (SDS-PAGE) and electro-transferred to a nitrocellulose membrane. Membranes were blocked for 1 hour at room temperature in 5% (w/v) BSA in Tris-buffered saline and 0.1% (v/v) Tween 20 (TBST) and then incubated overnight at 4°C with anti-ARC antibodies (1:500). After extensive washing with TBST, the membranes were incubated for 1 h at room temperature with the appropriate horseradish peroxidase (HRP)-conjugated secondary antibodies (1:5000, anti-mouse). After excessive washing with TBST, bound antibodies were visualized using ECL substrate for HRP and captured on ImageQuant LAS 500 imager system. Membranes were then stripped for 30 min at 50°C in buffer containing 100 mM 2-Mercaptoethanol, 2% (w/v) SDS and 62.5 mM Tris-HCl, pH 6.7, followed by extensive washing in TBST before re-blocking and re-probing with GAPDH-specific antibodies. The optical densities of the relevant immunoreactive bands were quantified using ImageLab software (version 4.1, Bio-Rad). The optical density values of the ARC protein immunoreactive bands were normalized to those of GAPDH. Results are expressed as a percentage of the values obtained with a control group.

### Quantitative reverse transcriptase polymerase chain reaction (qRT-PCR

Following brain dissection, tissue samples were immediately snap-frozen in liquid nitrogen and stored at −80°C until use. Frozen tissues were mechanically homogenized in TRIzol reagent and total RNA was isolated from each sample according to the manufacturer’s recommended protocol. mRNA was reverse transcribed to cDNA using the Reverse Transcription System and RevertAid kit. Plates (96 wells) were prepared for SYBR Green cDNA analysis using Fast SYBR Master Mix. Samples were analyzed in triplicate/duplicate with a Real-Time PCR System (StepOnePlus, Applied Biosystems), and quantified against an internal control gene *Gapdh*. We used the following reaction primers sequences: *Arc* forward, 5′-ACCGGGGGTCACTAAGTATGG -3′; reverse, 5′-CATTCTCCTGGCTCTGTAGGC -3′; *Egr1*(*Zif268*) forward, 5′-TGAGCACCTGACCACAGAGTC -3′; reverse, 5′-TAACTCGTCTCCACCATCGC-3′; *Bdnf IV* forward, 5′-GCAGCTGCCTTGATGTTTAC -3′; reverse, 5′-CCGTGGACGTTTACTTCTTTC -3′; *Gapdh* forward, 5′-CCAGAACATCATCCCTGC-3′; reverse, 5′-GGAAGGCCATGCCAGTGAGC-3′. Thermal cycling was initiated with incubation at 95°C for 20 s (for SYBR Green activation), followed by 40 cycles of PCR with the following conditions: Heating at 95°C for 3 s and then 30 s at 60°C. Relative quantification was calculated using the ΔΔCt method. The expression of target genes was normalized to that of *Gapdh*, and is expressed as percentage of the control group.

### RNA-seq library preparation

Frozen brain tissue was re-suspended in 0.6 mL TRI Reagent (Sigma T9424) and Dounce-homogenized for 40 strokes using tight pestle. RNA was purified using phenol-chloroform extraction and isopropanol precipitation and quality was assessed on an agarose gel. A 2 µg aliquot of total RNA from each sample was depleted of ribosomal RNAs using a Ribominus Eukaryote Kit v2 (Thermo Fisher Scientific, A15020) and processed using a previously published library preparation protocol^72^. Primers and adaptors used are listed in Table S1. The cDNA libraries were amplified using primers that carry Illumina indices, pooled, and 250-500 bp DNA fragments were isolated by agarose gel purification. The libraries were subjected to single-end 50-bp sequencing using the Illumina HiSeq 2000 platform. We utilized 2 to 3 biological replicates for each condition.

## Data analysis

Place preference was assessed as the percentage of time spent in the alcohol/sucrose-paired compartment, relative to the total test time. Mice that spent >75% of time in either of the compartments during Baseline test were excluded from the study (9 out of 267). CPP establishment was confirmed by comparing CPP scores between the baseline and CPP test 1, and was analyzed by a paired t-test. Since the expression of CPP (i.e., memory formation) is required for memory retrieval, data from mice that did not show CPP (a minimum 5% increase in preference between baseline and Test 1) were excluded from the experiment (48 out of 258).

Alcohol-CPP following interference with memory reconsolidation was analyzed by a mixed-model ANOVA with a between-subjects factor of Treatment (Actinomycin D or ODN) and a within-subjects factor of Test (CPP test 1, CPP test 2). Significant interactions were analyzed by a Student–Newman–Keuls post-hoc test. The densities of Western blot immunoreactive ARC protein levels were normalized to those GAPDH and analyzed by a one-way ANOVA with a between-subjects factor of Treatment, followed by Dunnett’s post-hoc test. qRT-PCR mRNA expression data were normalized to *Gapdh*, and expressed as percentage of expression of a control group. Data from all genes for each brain region were analyzed by a one-way MANOVA with a between-subjects factor of Treatment, followed by Dunnett’s test for each gene, for comparison to the data obtained from a control (No Retrieval) group.

For RNAseq, after confirming the quality of sequencing data by FastQC, reads were mapped to the mm9 reference genome using Bowtie2^73^ and annotated with Tophat2^74^. Reads mapped to genes were quantified by FeatureCounts^75^. We excluded *Rn45s, Lars2, Rn4*.*5s, Cdk8, Zc3h7a* and the mitochondrial chromosome to avoid counts of over-amplified genes that could skew library normalization. DESeq2^44^ was used to identify differentially expressed genes (DEG) in the Retrieval group, compared to the No Retrieval control group, with a significance cutoff adjusted p-value (p^(adj)^< 0.05).

## Data availability

RNA-seq data are available at NCBI Gene Expression Omnibus at https://www.ncbi.nlm.nih.gov/geo/query/acc.cgi (GSE205586).

## Acknowledgement

This work was supported by funds from the Israel Science Foundation (ISF) grants 968-13 and 1916-13 (SB), the Zuckerman STEM Leadership Program (KG), National Science Foundation Graduate Research Fellowship Program, DGE #1256260 (PMG), and National Institute of Health grants NS089896, NS125449, MH127485, and NS116008 (SI).

## Author contributions

KG and SB designed the research, KG and PG conducted the experiments. KG, PG, SI and SB analyzed the data and wrote the manuscript.

## Figure legends

**Table S1.**
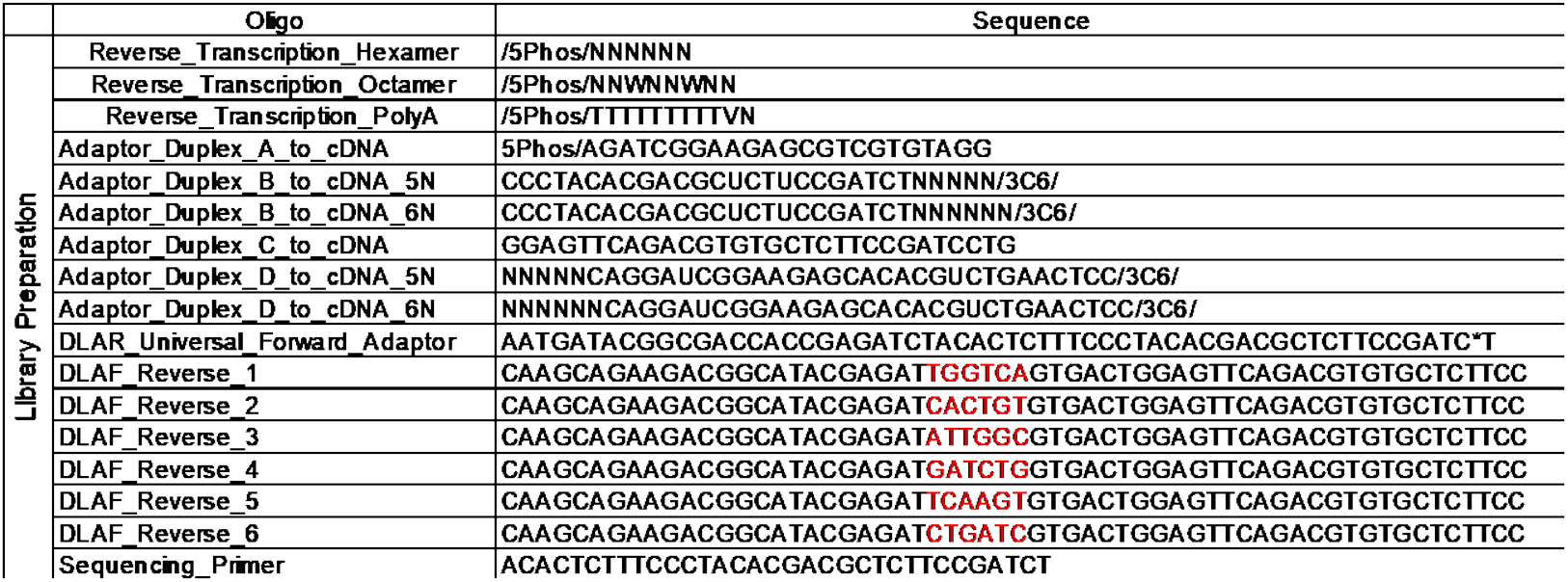
Primers and adaptors utilized in RNA-seq. All oligonucleotides are written in the 5’ to 3’direction; *indicates a phosphorothioate bond; /5Phos/ indicates 5’ phosphorylation; /3C6/ indicates a 3’ hexanediol; red highlight indicates the reverse complement of the 6 bp-long Illumina sequencing index.

**Table S2.**
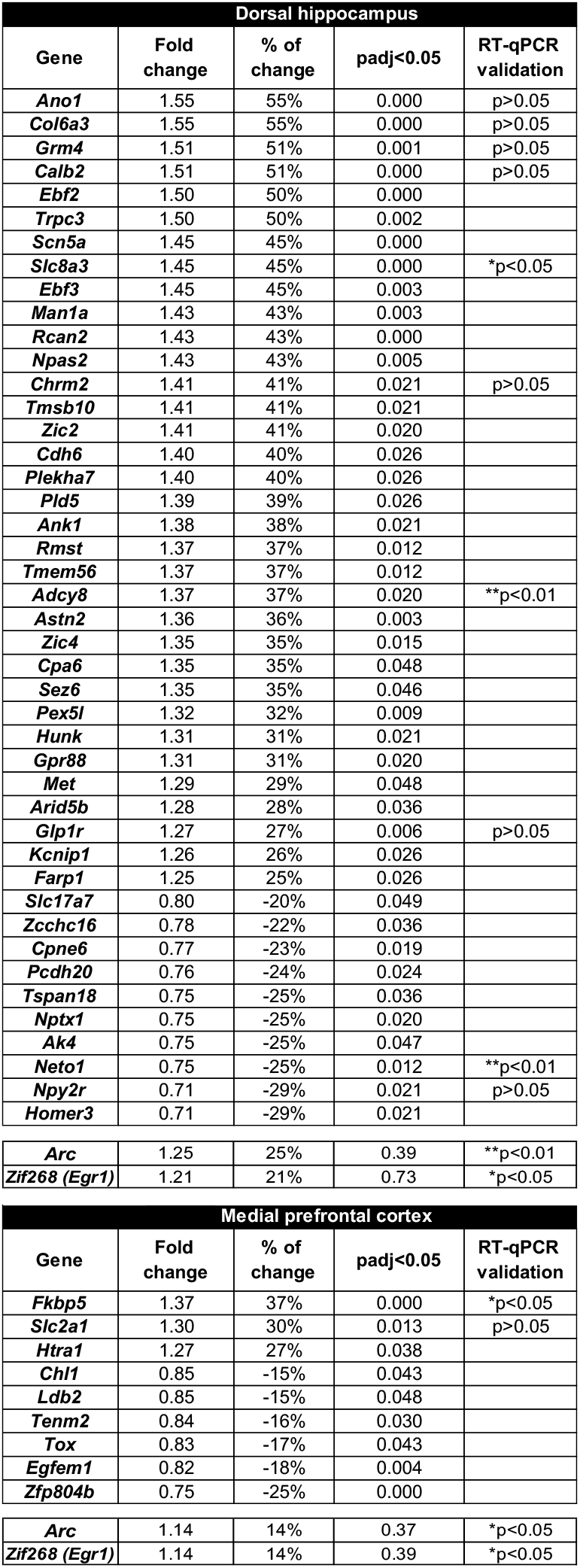
List of differentially expressed genes in the DH and mPFC following alcohol memory retrieval. The table contains estimates for the fold change and percent of change in the expression of differentially expressed genes in the Retrieval group, as compared to No Retrieval group. The table also lists differentially expressed genes validated by qRT-PCR analysis.

| Dorsal hippocampus |  |  |  |  |
| --- | --- | --- | --- | --- |
| Gene | Fold change | % of change | padj<0.05 | RT-qPCR validation |
| *Ano1* | 1.55 | 55% | 0.000 | p>0.05 |
| *Col6a3* | 1.55 | 55% | 0.000 | p>0.05 |
| *Grm4* | 1.51 | 51% | 0.001 | p>0.05 |
| *Calb2* | 1.51 | 51% | 0.000 | p>0.05 |
| *Ebf2* | 1.50 | 50% | 0.000 |  |
| *Trpc3* | 1.50 | 50% | 0.002 |  |
| *Scn5a* | 1.45 | 45% | 0.000 |  |
| *Slc8a3* | 1.45 | 45% | 0.000 | *p<0.05 |
| *Ebf3* | 1.45 | 45% | 0.003 |  |
| *Man1a* | 1.43 | 43% | 0.003 |  |
| *Rcan2* | 1.43 | 43% | 0.000 |  |
| *Npas2* | 1.43 | 43% | 0.005 |  |
| *Chrm2* | 1.41 | 41% | 0.021 | p>0.05 |
| *Tmsb10* | 1.41 | 41% | 0.021 |  |
| *Zic2* | 1.41 | 41% | 0.020 |  |
| *Cdh6* | 1.40 | 40% | 0.026 |  |
| *Plekha7* | 1.40 | 40% | 0.026 |  |
| *Pld5* | 1.39 | 39% | 0.026 |  |
| *Ank1* | 1.38 | 38% | 0.021 |  |
| *Rmst* | 1.37 | 37% | 0.012 |  |
| *Tmem56* | 1.37 | 37% | 0.012 |  |
| *Adcy8* | 1.37 | 37% | 0.020 | **p<0.01 |
| *Astn2* | 1.36 | 36% | 0.003 |  |
| *Zic4* | 1.35 | 35% | 0.015 |  |
| *Cpa6* | 1.35 | 35% | 0.048 |  |
| *Sez6* | 1.35 | 35% | 0.046 |  |
| *Pex5l* | 1.32 | 32% | 0.009 |  |
| *Hunk* | 1.31 | 31% | 0.021 |  |
| *Gpr88* | 1.31 | 31% | 0.020 |  |
| *Met* | 1.29 | 29% | 0.048 |  |
| *Arid5b* | 1.28 | 28% | 0.036 |  |
| *Glp1r* | 1.27 | 27% | 0.006 | p>0.05 |
| *Kcnip1* | 1.26 | 26% | 0.026 |  |
| *Farp1* | 1.25 | 25% | 0.026 |  |
| *Slc17a7* | 0.80 | -20% | 0.049 |  |
| *Zcchc16* | 0.78 | -22% | 0.036 |  |
| *Cpne6* | 0.77 | -23% | 0.019 |  |
| *Pcdh20* | 0.76 | -24% | 0.024 |  |
| *Tspan18* | 0.75 | -25% | 0.036 |  |
| *Nptx1* | 0.75 | -25% | 0.020 |  |
| *Ak4* | 0.75 | -25% | 0.047 |  |
| *Neto1* | 0.75 | -25% | 0.012 | **p<0.01 |
| *Npy2r* | 0.71 | -29% | 0.021 | p>0.05 |
| *Homer3* | 0.71 | -29% | 0.021 |  |
| *Arc* | 1.25 | 25% | 0.39 | **p<0.01 |
| *Zif268 (Egr1)* | 1.21 | 21% | 0.73 | *p<0.05 |

**Table S2. List of differentially expressed genes in the DH and mPFC following alcohol memory retrieval.**
| Medial prefrontal cortex |  |  |  |  |
| --- | --- | --- | --- | --- |
| Gene | Fold change | % of change | padj<0.05 | RT-qPCR validation |
| *Fkbp5* | 1.37 | 37% | 0.000 | *p<0.05 |
| *Slc2a1* | 1.30 | 30% | 0.013 | p>0.05 |
| *Htra1* | 1.27 | 27% | 0.038 |  |
| *Chl1* | 0.85 | -15% | 0.043 |  |
| *Ldb2* | 0.85 | -15% | 0.048 |  |
| *Tenm2* | 0.84 | -16% | 0.030 |  |
| *Tox* | 0.83 | -17% | 0.043 |  |
| *Egfem1* | 0.82 | -18% | 0.004 |  |
| *Zfp804b* | 0.75 | -25% | 0.000 |  |
| *Arc* | 1.14 | 14% | 0.37 | *p<0.05 |
| *Zif268 (Egr1)* | 1.14 | 14% | 0.39 | *p<0.05 |

**Figure S1.**
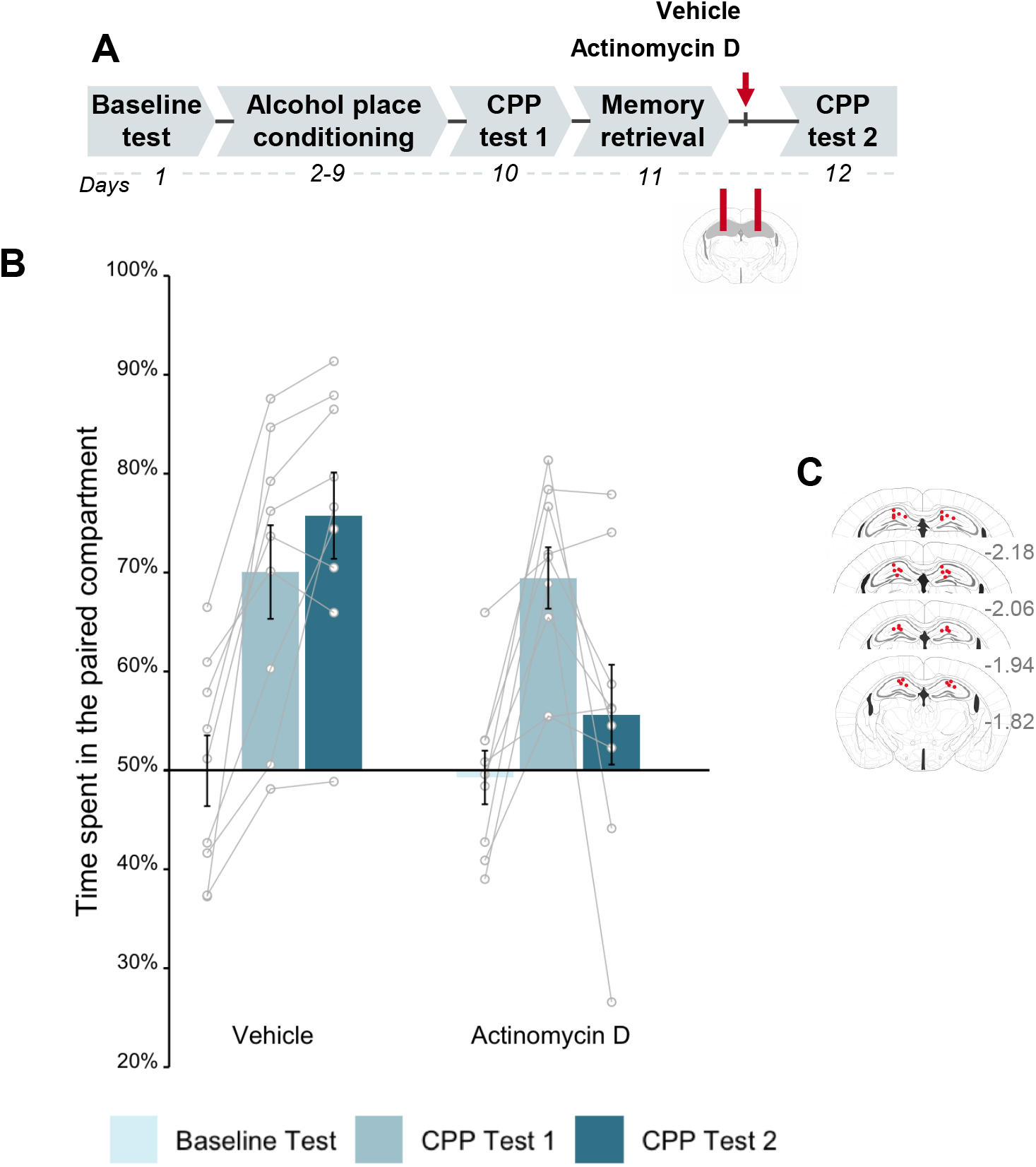
Inhibition of transcription in the dorsal hippocampus after alcohol memory retrieval disrupts the expression of alcohol-CPP. **A**. Schematic illustration of the experimental design and timeline. Actinomycin D (4 µg/µl) was bilaterally infused into the dorsal hippocampus of mice immediately following the retrieval of alcohol memories. **B**. Place preference scores, expressed as means ±S.E.M. of the percent of time spent in the alcohol-paired compartment. **C**. Locations of cannulas. *p<0.05; n=9 per group.

**Figure S2.**
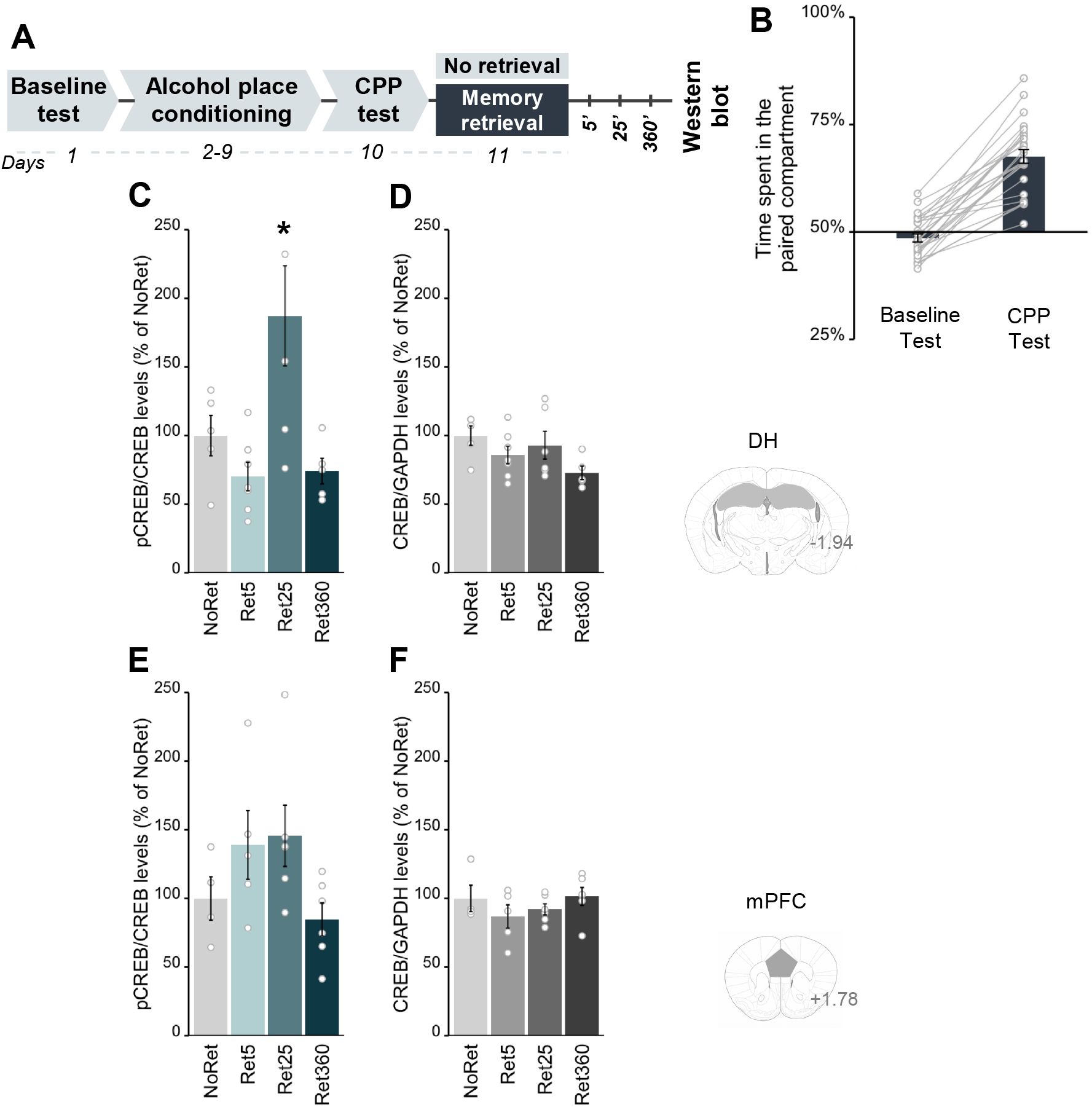
Alcohol memory retrieval induces CREB phosphorylation in the DH and mPFC. **A**. Schematic illustration of the experimental design and timeline. **B**. Place preference scores, expressed as means ±S.E.M.. of the percent of time spent in the alcohol-paired compartment; **C-F**. Protein levels, expressed as means ±S.E.M. of the percent of change from the control group (No Retrieval). Levels of pCREB were normalized to of tCREB in the DH. The levels of pCREB in the DH were increased 25 min after alcohol-memory retrieval (one-way ANOVA; Time (F(3,18)=6.07, p<0.05); post hoc: NoRet vs Ret25’ (p<0.05)) (**C**) or mPFC (**E**). Levels of tCREB were normalized to GAPDH in the DH (**D**) or mPFC (**F**).; *p<0.05; n=5-6 per group.

**Figure S3.**
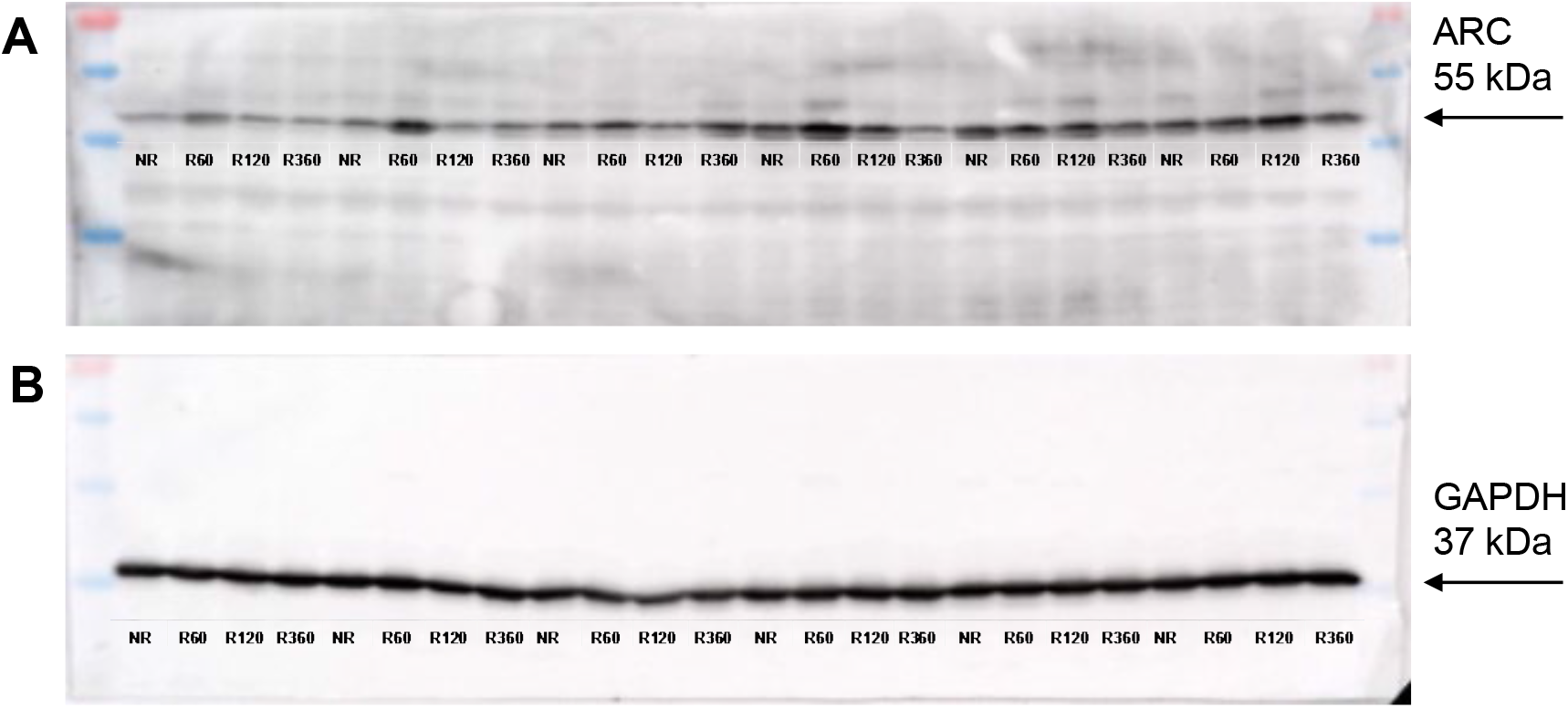
ARC (**A**) and GAPDH (**B**) protein expression in the DH following alcohol-memory retrieval, as revealed by Western blot.

**Figure S4.**
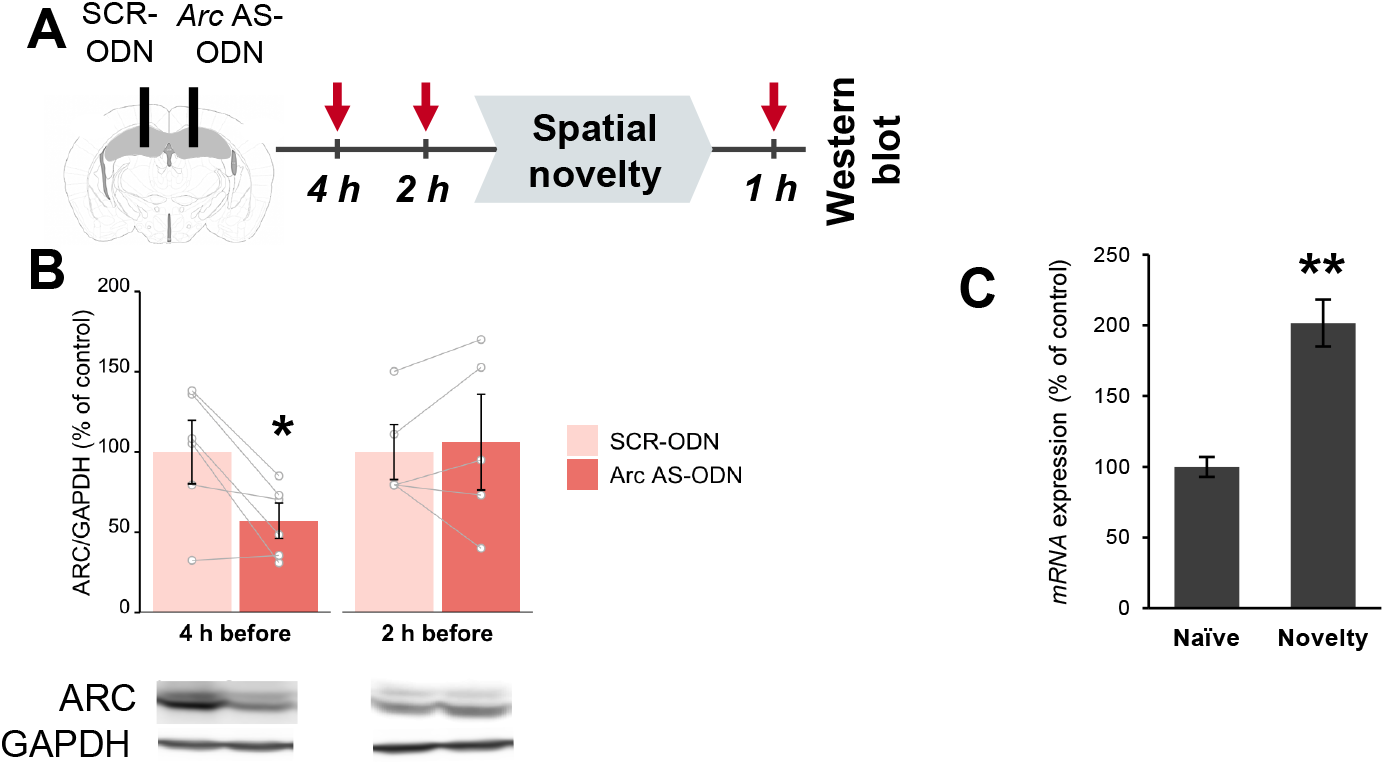
Validation of *Arc* AS-ODN. **A**. Schematic representation of the validation of the ability of *Arc* AS-ODN to downregulate ARC protein expression in the DH. To this end, we infused *Arc* AS-ODN (2 nmol/µl, 0.5 µl) into one hemisphere, and an equivalent amount of a non-specific SCR-ODN into the other hemisphere (sides were counter-balanced). Two or four hours later, we allowed the mice to explore an unfamiliar environment (CPP compartment) for 5 min to induce novelty-dependent ARC expression. One hour later (3 or 5 h after ODN infusion), brain tissues were collected, and the levels of ARC protein were assessed by Western blot. **B**. Protein levels, expressed as means ±S.E.M. of the percent of change from the control hemisphere (SCR-ODN). Levels of ARC were normalized to GAPDH. We found that *Arc* AS-ODN produced a 40% decrease in novelty-induced ARC protein levels in the DH, when administered 4 h (t(5)=-3.29, p<0.05), but not 2 h (t(5)=0.44, p>0.05), prior to the novelty presentation, as compared to the SCR-ODN control. These results show that infusion of *Arc* AS-ODN into the DH downregulated ARC protein levels 5 h, but not 3 h, after infusion. **C**. mRNA levels, normalized to *Gapdh*, as the percent of change from the control group (naive). qRT-PCR analysis revealed a rapid 2-fold change in *Arc* expression in the DH in mice 30 min after a 5 min novelty exploration period (p<0.01). *p<0.05; **p<0.01; n=4-6 per group.

**Figure S5.**
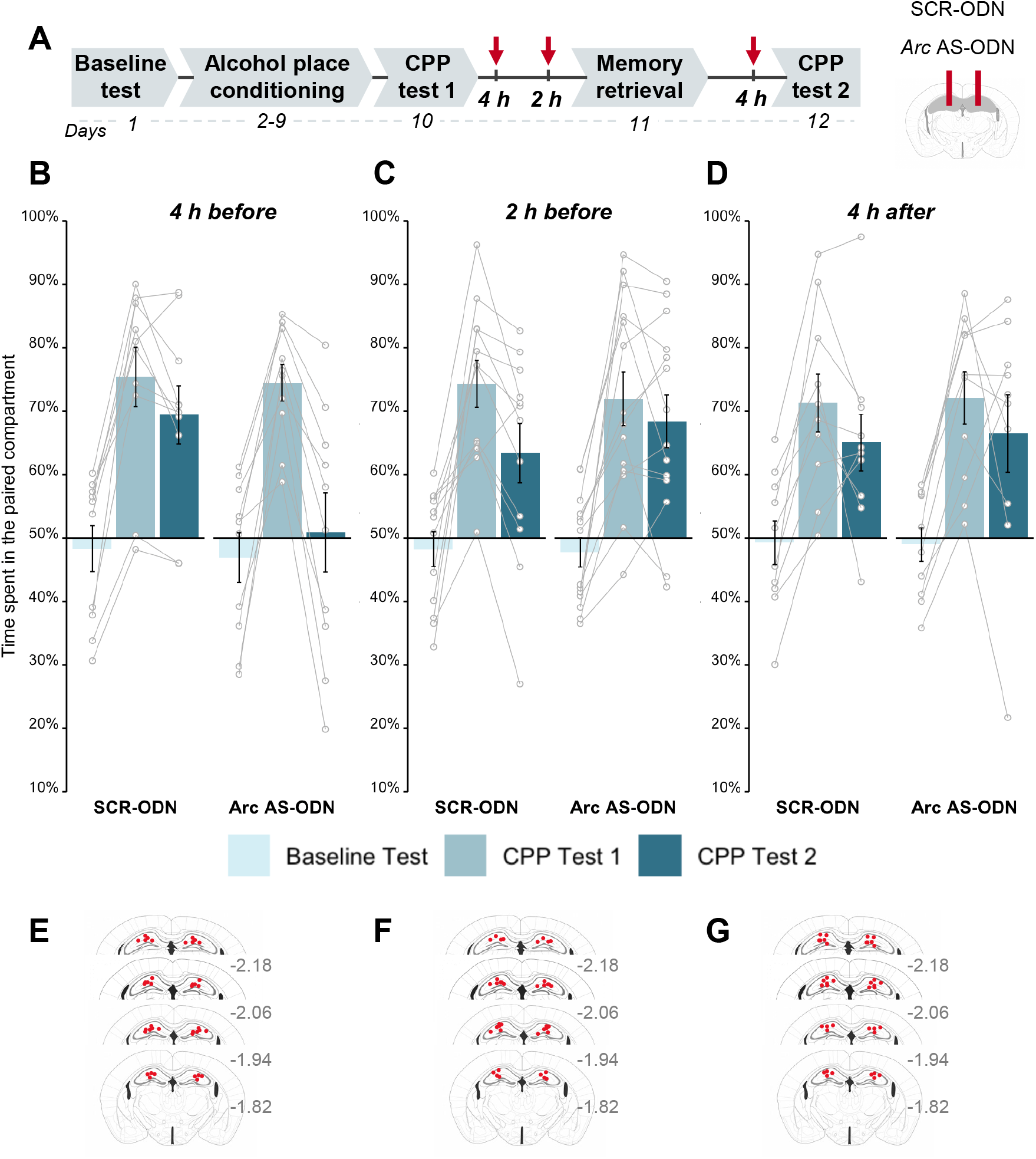
Downregulation of ARC protein in the dorsal hippocampus shortly after alcohol memory retrieval disrupts the expression of alcohol-CPP. **A**. Schematic illustration of the experimental design and timeline. Antisense oligodeoxynucleotides directed against *Arc* mRNA (*Arc* AS-ODN) or non-specific scrambled oligodeoxynucleotides (SCR-ODN) were infused into the dorsal hippocampus (DH) of mice at the indicated time points. **B-D**. Place preference scores, expressed as means ±S.E.M. of the percent of time spent in the alcohol-paired compartment. Infusion of *Arc* AS-ODN disrupted the expression of alcohol-CPP only when infused 4 h **(B)**, but not 2 h before memory retrieval **(C)** or 4 h after memory retrieval **(D)**, as compared with SCR-ODN-treated controls. **E-G**. Locations of cannulas. *p<0.05; n=10-12 per group.

